# Syndecan-1 as specific cerebrospinal fluid biomarker of multiple sclerosis

**DOI:** 10.1101/2023.05.10.540204

**Authors:** Geoffrey Hinsinger, Lucile Du Trieu de Terdonck, Serge Urbach, Nicolas Salvetat, Manon Rival, Manon Galoppin, Chantal Ripoll, Renaud Cezar, Sabine Laurent-Chabalier, Christophe Demattei, Hanane Agherbi, Giovanni Castelnovo, Sylvain Lehmann, Valérie Rigau, Philippe Marin, Eric Thouvenot

**Author notes:** Correspondance to: Dr Thouvenot, Institut de Génomique Fonctionnelle, 141 rue de la Cardonille, F-34094 Montpellier Cedex 5, France. Phone n°, Fax n°. Authors contributed equally to this work (Geoffrey Hinsinger, Lucile Du Trieu de Terdonck). **Abbreviations**: CECR1: Cat eye syndrome critical region protein 1; CHI3L1: Chitinase 3-like protein 1; CHI3L2: Chitinase 3-like protein 2; CHIT1: chitotriosidase-1; CIS: clinically isolated syndrome; CSF, cerebrospinal fluid; CTRL: symptomatic control; EDSS: Expanded Disability Status Scale; IGKC: immunoglobulin kappa chain region C; ION: isolated optic neuritis; INDC: inflammatory neurological disease control; MS: multiple sclerosis; NAWM: normal appearing white matter; NGAL: neutrophil gelatinase-associated lipocalin; NINDC: non-inflammatory neurological disease controls; OCBs: oligoclonal bands; PINDC: peripheral inflammatory neurological disease control; PPMS: primary progressive multiple sclerosis; RRMS: relapsing-remitting multiple sclerosis; SDC1: syndecan-1; WBCs: white blood cells.

## Abstract

Multiple sclerosis (MS) is an inflammatory demyelinating disease often characterized by remission and relapse periods occurring at irregular intervals after an initial attack (clinically isolated syndrome) and followed by a gradual progression of disability. Clinical symptoms, magnetic resonance imaging and abnormalities in cerebrospinal fluid (CSF) immunoglobulin profile allow diagnosis with a good sensitivity. However, current biomarkers lack specificity or have poor individual prognostic value. To identify novel candidate biomarkers of MS, we analysed 1) the CSF proteome from symptomatic controls and patients with clinically isolated syndrome or remitting-relapsing multiple sclerosis (n=40), and 2) changes in oligodendrocyte secretome upon proinflammatory or pro-apoptotic treatment. Proteins exhibiting differences in abundance in both studies were combined with previously described MS biomarkers to build a list of 87 proteins that were quantified by parallel reaction monitoring (PRM) in CSF samples from a new cohort comprising symptomatic controls and MS patients at different disease stages (n=60). The eleven proteins that passed this qualification step were subjected to a new PRM assay from a larger cohort (n=158) comprising patients with MS at different disease stages or with other inflammatory or non-inflammatory neurological disorders. Collectively, these studies identified a biomarker signature of MS that might improve MS diagnosis and prognosis. These include the oligodendrocyte precursor cell proteoglycan Syndecan-1, which was more efficient than previously described biomarkers to discriminate MS from other inflammatory and non-inflammatory neurological disorders.

## Introduction

Multiple sclerosis (MS) is an inflammatory autoimmune disease of the central nervous system (CNS) that causes damage to myelin along with axons and ultimately leads to neurodegeneration. MS diagnosis mainly relies on clinical symptoms and the presence of CNS lesions detected by magnetic resonance imaging (MRI). Diagnostic criteria for MS recently evolved to better predict disease activity in patients experiencing a clinically isolated syndrome (CIS) for early and personalized therapy (1–3). However, dissemination in time, assessed by the presence of active lesions in MRI or the presence of oligoclonal bands (OCBs) in cerebrospinal fluid (CSF) only partially predict relapses and disability progression. There is still a need for biomarkers of disease activity as well as MS differential diagnosis and important efforts are being made to identify specific biomarkers of the disease. The CSF represents a fluid of choice to investigate MS pathophysiology and identify specific biomarkers. Several proteomic studies have compared the proteome of CSF samples from either RRMS patients and controls (including healthy and/or symptomatic controls), or MS patients at different disease stages (CIS, relapsing-remitting multiple sclerosis (RRMS) and primary progressive multiple sclerosis (PPMS). They identified chitinase-like proteins (CHI3L1 and CHI3L2) and chitotriosidase (CHIT1) as potential biomarkers of MS, in comparison with symptomatic controls or isolated optic neuritis (4–7). However, their CSF levels are also elevated in other neurological inflammatory and non-inflammatory conditions (8,9), and cannot be used as biomarkers for differential diagnosis in addition to CSF immunoelectrophoresis and MRI. More recently, neurofilament-light (NfL) and heavy (NfH) chains were shown to be elevated both in the CSF and serum of MS patients (10,11). Again, these biomarkers of neuronal/axonal damage are not specific for MS, as they are also increased in neurodegenerative and neuroinfectious diseases (12).

Accordingly, MS diagnosis still relies on the association of a suggestive clinical presentation, the presence of typical MRI lesions of the CNS, and the ancient CSF biomarker OCBs. Over the past years, the kFLC index emerged as a more specific candidate biomarker of MS that reflects intrathecal synthesis of immunoglobulins, as quantitative substitute of OCBs (13,14).

However, it is not widely used in clinical practice, and cut-off values discriminating MS and other disorders differ from one study to one another (15–18). Differential diagnosis of MS remains challenging given the common feature of MS and other neurological inflammatory and non-inflammatory conditions (19).

To address this issue, we combined for the first time quantitative proteomic analysis of CSF samples from symptomatic controls and patients with clinically isolated syndrome or remitting-relapsing multiple sclerosis (n=40) with analysis of changes in the secretome of primary cultured rat oligodendrocyte precursor cells (OPCs) in response a proinflammatory or a proapoptotic treatment to mimic the main pathological features of the disease *in vitro*. Proteins exhibiting differences in abundances between the different groups in both studies were combined with previously identified biomarkers of MS not detected in our study to build a list of 87 proteins that were quantified by parallel reaction monitoring (PRM) in CSF samples from the discovery cohort and a new cohort comprising symptomatic controls and MS patients at different disease stages (n=60). The eleven proteins that passed this qualification step were then subjected to a second PRM assay from a larger cohort (n=158) comprising patients with MS at different disease stages or with other inflammatory or non-inflammatory neurological disorders. This second PRM assay revealed biomarker signatures that segregate between MS and other inflammatory or non-inflammatory neurological disorders and identified the cell surface proteoglycan Syndecan-1 (SDC1) as a novel specific CSF biomarker of MS.

## Materials and Methods

### Patients

Symptomatic controls (CTRLs) and patients with CIS, RRMS, PPMS, inflammatory neurological disease control (INDC), peripheral inflammatory neurological disease control (PINDC) and non-inflammatory neurological disease controls (NINDC) were recruited at Montpellier and Nîmes multiple sclerosis centres and prospectively followed for at least two years. All enrolled patients have signed informed consent for CSF biobanking after lumbar puncture prescribed for investigation of neurological disorders (biobank registered under n° DC-2008-417). The procedures were authorized by the Agence Nationale de Sécurité des Médicaments et des produits de santé (ANSM, n°ID RCB 2008-A01199-46) on September 12, 2008 and approved by the Comité de Protection des Personnes Sud Méditerranée IV (ethics committee) on February 10, 2009. CTRLs included patients explored for headaches, paresthesia, visual disturbance, vertigo or dizziness (20). All presented a normal clinical examination, normal biological tests, normal CSF (including absence of OCBs and normal IgG index), and normal brain and spinal MRI. Patients with isolated optic neuritis (ION) had no abnormal brain or spinal cord lesion and normal CSF. A schematic definition of CIS, RRMS, PPMS and controls is provided on Supplementary Fig. 1. CIS patients were defined as patients experiencing a first neurological attack typical of an MS relapse with MRI lesions fulfilling the Swanton criteria for dissemination in space (DIS), including patients fulfilling Barkhof-Tintore criteria for DIS and/or the 2017 McDonald’s criteria for MS. CIS patients were followed clinically and by MRI in a prospective manner from the first attack without receiving any disease-modifying treatment before conversion to RRMS. Conversion of CIS to RRMS was assessed using the McDonald criteria revised in 2005, after a 2^nd^ relapse or the apparition of new MRI lesions on follow-up scans (1). CIS patients with fast-conversion to RRMS (< 1 year after the CIS FC-CIS) and patients with slow-conversion to RRMS (> 2 years after a CIS, SC-CIS) were included in the study (Supplementary Fig. 1). RRMS patients had lumbar puncture at the time of the first relapse. PPMS was defined according to the McDonald criteria revised in 2005 (Supplementary Fig. 1). In the verification cohort, INDCs included patients with encephalitis (n=4), isolated myelitis (n=3), acute demyelinating encephalomyelitis (ADEM, n=2), chronic lymphocytic inflammation with pontine perivascular enhancement responsive to steroids (CLIPPERS, n=1), aseptic meningitis (n=1), cerebral toxoplasmosis (n=1) and sinus inflammation related optic neuritis (n=1); PINDCs included patients with acute inflammatory demyelinating polyneuropathy (AIDP, n=4), chronic inflammatory demyelinating polyneuropathy (CIDP, n=4), polyneuritis (n=2), plexopathy (n=2), and multiple mononeuropathy (n=1); NINDCs included patients with transient ischemic attack (TIA) with small vessel disease (n=3), frontotemporal dementia (n=2), stroke (n=1), generalized dystonia (n=1), adrenoleukodystrophy (n=1), cerebellar ataxia (n=1), hydrocephaly (n=1), syringomyelia (n=1), sacral plexus compression (n=1), and metabolic encephalitis (n=1) (20). Demographics, CSF and MRI characteristics of patients used in the discovery, qualification and verification steps of proteomic analysis are described in Table 1 and Table 2.

**Table 1:**
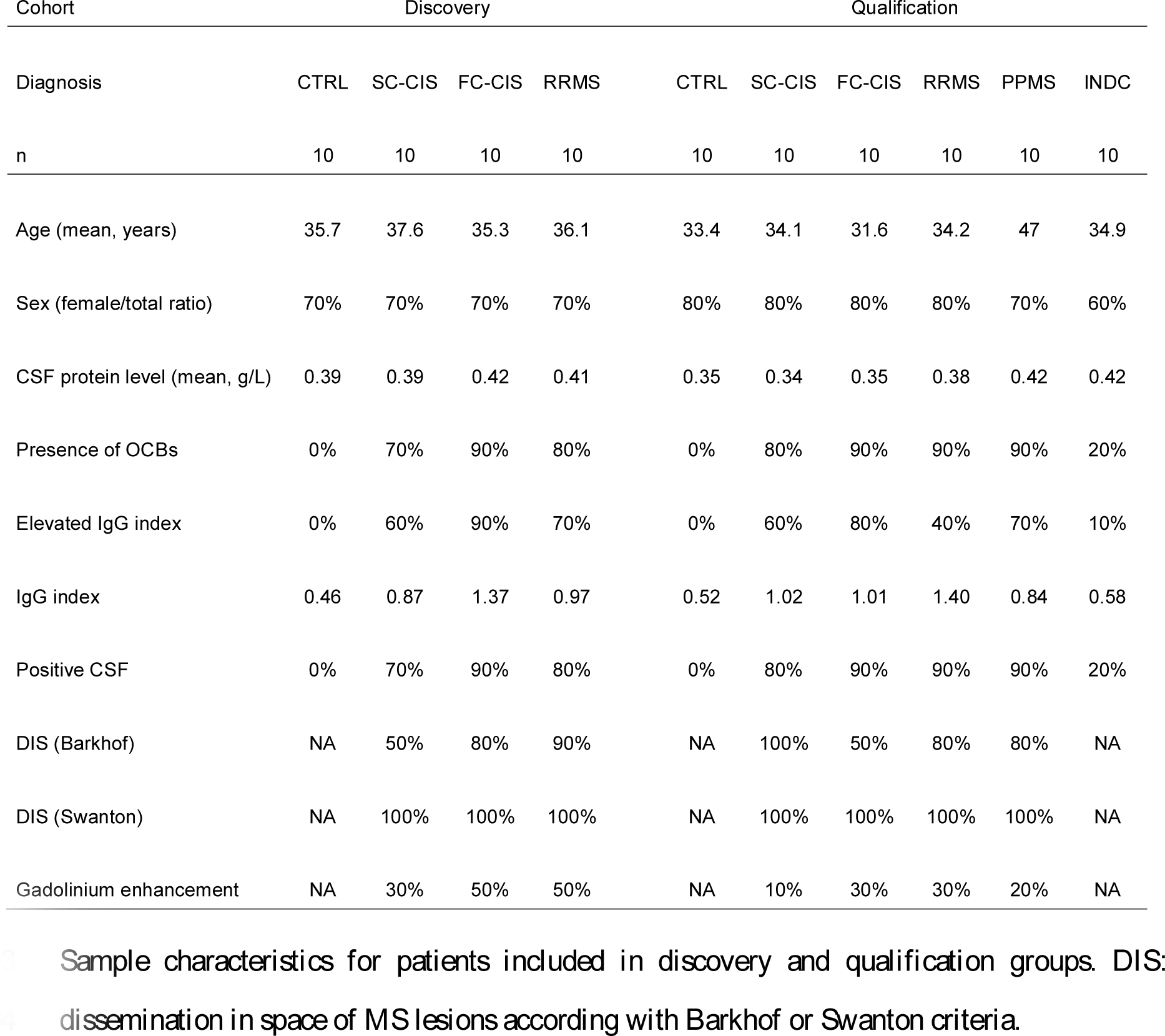
Characteristics of the patients included in the discovery and qualification cohorts.

| Cohort | Discovery |  |  |  | Qualification |  |  |  |  |  |
| --- | --- | --- | --- | --- | --- | --- | --- | --- | --- | --- |
| Diagnosis | CTRL | SC-CIS | FC-CIS | RRMS | CTRL | SC-CIS | FC-CIS | RRMS | PPMS | INDC |
| n | 10 | 10 | 10 | 10 | 10 | 10 | 10 | 10 | 10 | 10 |
| Age (mean, years) | 35.7 | 37.6 | 35.3 | 36.1 | 33.4 | 34.1 | 31.6 | 34.2 | 47 | 34.9 |
| Sex (female/total ratio) | 70% | 70% | 70% | 70% | 80% | 80% | 80% | 80% | 70% | 60% |
| CSF protein level (mean, g/L) | 0.39 | 0.39 | 0.42 | 0.41 | 0.35 | 0.34 | 0.35 | 0.38 | 0.42 | 0.42 |
| Presence of OCBs | 0% | 70% | 90% | 80% | 0% | 80% | 90% | 90% | 90% | 20% |
| Elevated IgG index | 0% | 60% | 90% | 70% | 0% | 60% | 80% | 40% | 70% | 10% |
| IgG index | 0.46 | 0.87 | 1.37 | 0.97 | 0.52 | 1.02 | 1.01 | 1.40 | 0.84 | 0.58 |
| Positive CSF | 0% | 70% | 90% | 80% | 0% | 80% | 90% | 90% | 90% | 20% |
| DIS (Barkhof) | NA | 50% | 80% | 90% | NA | 100% | 50% | 80% | 80% | NA |
| DIS (Swanton) | NA | 100% | 100% | 100% | NA | 100% | 100% | 100% | 100% | NA |
| Gadolinium enhancement | NA | 30% | 50% | 50% | NA | 10% | 30% | 30% | 20% | NA |

Sample characteristics for patients included in discovery and qualification groups. DIS: dissemination in space of MS lesions according with Barkhof or Swanton criteria.

**Table 2:** Characteristics of the patients included in the verification cohort.

| Diagnosis | CTRL | SC-CIS | FC-CIS | RRMS | PPMS | INDC | ION | NINDC | PINDC |
| --- | --- | --- | --- | --- | --- | --- | --- | --- | --- |
| n | 30 | 15 | 15 | 30 | 14 | 13 | 15 | 13 | 13 |
| Age (mean, years) | 38.3 | 35.2 | 33.3 | 38.2 | 46.6 | 46.4 | 31.8 | 40.8 | 56.2 |
| Sex (female/total ratio) | 80% | 73% | 93% | 77% | 43% | 23% | 87% | 46% | 54% |
| CSF protein level (mean, g/L) | 0.36 | 0.38 | 0.35 | 0.39 | 0.51 | 0.39 | 0.36 | 0.33 | 0.41 |
| Presence of OCBs | 0% | 80% | 93% | 90% | 79% | 15% | 7% | 0% | 15% |
| Elevated IgG index | 0% | 46% | 75% | 78% | 58% | 0% | 0% | 0% | 0% |
| IgG index | 0.48 | 0.68 | 0.86 | 1.12 | 0.84 | 0.47 | 0.47 | 0.45 | 0.48 |
| Positive CSF | 0% | 80% | 93% | 90% | 79% | 15% | 7% | 0% | 15% |
| DIS (Barkhof) | NA | 40% | 87% | 80% | 79% | NA | 0% | NA | NA |
| DIS (Swanton) | NA | 100% | 100% | 100% | 100% | NA | 0% | NA | NA |
| Gadolinium enhancement | NA | 27% | 47% | 47% | 14% | NA | 0% | NA | NA |

Sample characteristics for patients included the verification cohort. DIS = dissemination in space of white matter lesions according with Barkhof or Swanton criteria.

MS brain samples were obtained from a 50-year-old man who died during a fulminating MS relapse after Natalizumab withdrawal and presented extensive and active brain lesions characteristic of MS (21), a 47-year-old man with secondary progressive MS (SPMS) who died from a lung neoplasm, a 60-year-old woman with SPMS who died from an infection. Samples were rapidly (within 12 h after death) frozen in liquid nitrogen or fixed by immersion in 10% buffered formalin and processed into liquid paraffin for histological evaluation and immunohistochemistry (Centre des Collections Biologiques Hospitalières de Montpellier (CCBH-M), Collection tumorothèque, FINESS 340780477, F-34285 Montpellier, France; and Collection Sclérose en Plaques Nantes (PFS13-003), FINESS 440000289, F-44093, Nantes, France). Control brain samples were obtained from the right frontal lobe of a 62-year-old man who died after an occipital infarct and from the right frontal lobe of a 87-year-old man who died from pneumonia.

### CSF sample collection and preparation

CSF samples were collected using a 25G Whitacre-type atraumatic needle (ref 181.05, Vygon) in a 10 ml polypropylene tube (ref 62.610.201, Sarstedt) at the end of lumbar punctures (L3-L5). They were centrifuged at the latest 2 h after collection at 1,500 × *g* for 10 min at 4°C, according to the guidelines of the BioMS-eu network, except for sample transport and centrifugation at 4°C instead of room temperature (22). Aliquots (500 µl) were stored at - 80°C in 1.5 ml tubes (Protein LoBind 0030108.116, Eppendorf) until use. Patients with traumatic lumbar punctures (>500 red cells/mm^3^) were excluded.

### Quantitative proteomic analysis of CSF

#### Label-free quantitative proteomics

CSF samples (200 μl) were immunodepleted of the 20 most abundant plasma proteins (albumin, apolipoproteins A1, A2 and B, α-1-acid-glycoprotein, α-1-antitrypsin, α-2-macroglobulin, ceruloplasmin, complements C1q, C3 and C4, haptoglobin, fibrinogen, IgA, IgD, IgG, IgM, plasminogen, transferrin and transthyretin) using the ProteoPrep 20 plasma protein immunodepletion kit (Sigma), by performing two immunodepletion cycles, as previously described (23).

After reduction (with 20 mM dithiothreitol in presence of 6 M urea and 0.03% anionic acid labile surfactant (AALS) at 56°C for 15 min) and alkylation (55 mM iodoacetamide in presence of 6 M urea and 0.03% AALS at room temperature for 30 min), immunodepleted protein samples were successively digested with LysC (0.008 μg/μL, Wako) and trypsin (0.002 μg/μL, Gold, Promega), using a filter-aided sample preparation (FASP) procedure adapted from (24).

The resulting peptides were analyzed online by nano-flow HPLC-nanoelectrospray ionization using a Q-Exactive+ mass spectrometer (Thermo Fisher Scientific) coupled to a nano-LC system (U3000-RSLC, Thermo Fisher Scientific). Desalting and preconcentration of samples were performed on-line on a Pepmap® precolumn (0.3 × 10 mm; Thermo Fisher Scientific). A gradient consisting of 0–26% B in A for 120 min (A: 0.1% formic acid, 2% acetonitrile in water, and B: 0.1% formic acid in 80% acetonitrile), then 26-52% B for 20 min at 300 nl/min, was used to elute peptides from the capillary reverse-phase column (0.075 × 150 mm, Pepmap®, Thermo Fisher Scientific). Data were acquired using the Xcalibur software. A cycle of one full-scan mass spectrum (350–1,500 m/z) at a resolution of 70,000 (at 200 m/z), followed by 10 data-dependent MS/MS spectra (at a resolution of 17,500, isolation window 1.2 m/z) was repeated continuously throughout the nano-LC separation. Raw data were analysed using the MaxQuant software (version 1.4.1.2) (25) and the Andromeda search engine [http://coxdocs.org/doku.php?id=maxquant:andromeda:start] against the UniProtKB Reference proteome UP000005640 database for Homo sapiens (release 2013-07) and the contaminant database in MaxQuant. The following parameters were used: enzyme specificity set as Trypsin/P with a maximum of two missed cleavages, oxidation (M) and phosphorylation (STY) set as variable modifications and carbamidomethyl (C) as fixed modification, and a mass tolerance of 0.5 Da for fragment ions. The maximum false peptide and protein discovery rate was specified as 0.01 and minimum peptide length to 7. Relative protein quantification in the different CSF samples was performed using the label-free quantification (LFQ) algorithm [https://maxquant.net/maxquant/].

#### Statistical analysis of label-free quantitative data

Data were analyzed with the “R/Bioconductor” statistical open-source software (Gentleman, Carey et al. 2004). Before differential analysis, protein intensities were transformed in log2(X) and missing data was imputed by KNNimput approach (imputation R package). The differential intensity levels of proteins or peptides between groups were analyzed using different statistical tests Limma method (R package limma), SAM method (R package siggenes) and the most appropriate statistical test between Wilcoxon’s, Student’s or Welch’s test (control of normality and homoscedasticity hypothesis) were selected. With the multiple testing methodologies, it is important to adjust the p-value of each protein or peptide to control the False Discovery Rate (FDR). The Benjamini and Hochberg procedure (26) was applied on all statistical tests (Multtest R package). The fold change (Nfold) using the median has also been calculated and a value ±1.5 was considered as significant. The accuracy of each biomarker and its discriminatory power was evaluated using a receiver operating characteristic (ROC) analysis. ROC curves are the graphical visualisation of the reciprocal relation between the sensitivity (Se) and the specificity (Sp) of a test for various values (pROC R package).

For each significantly differential protein or peptide between two patient groups with one of the statistical tests used, a value was assigned according to Table 0. Thus, all proteins or peptides had a global score for all statistical tests between 0 and 4.5. Only spots with a score greater than or equal to 2 were included in the further analyses.

**Table 0.**
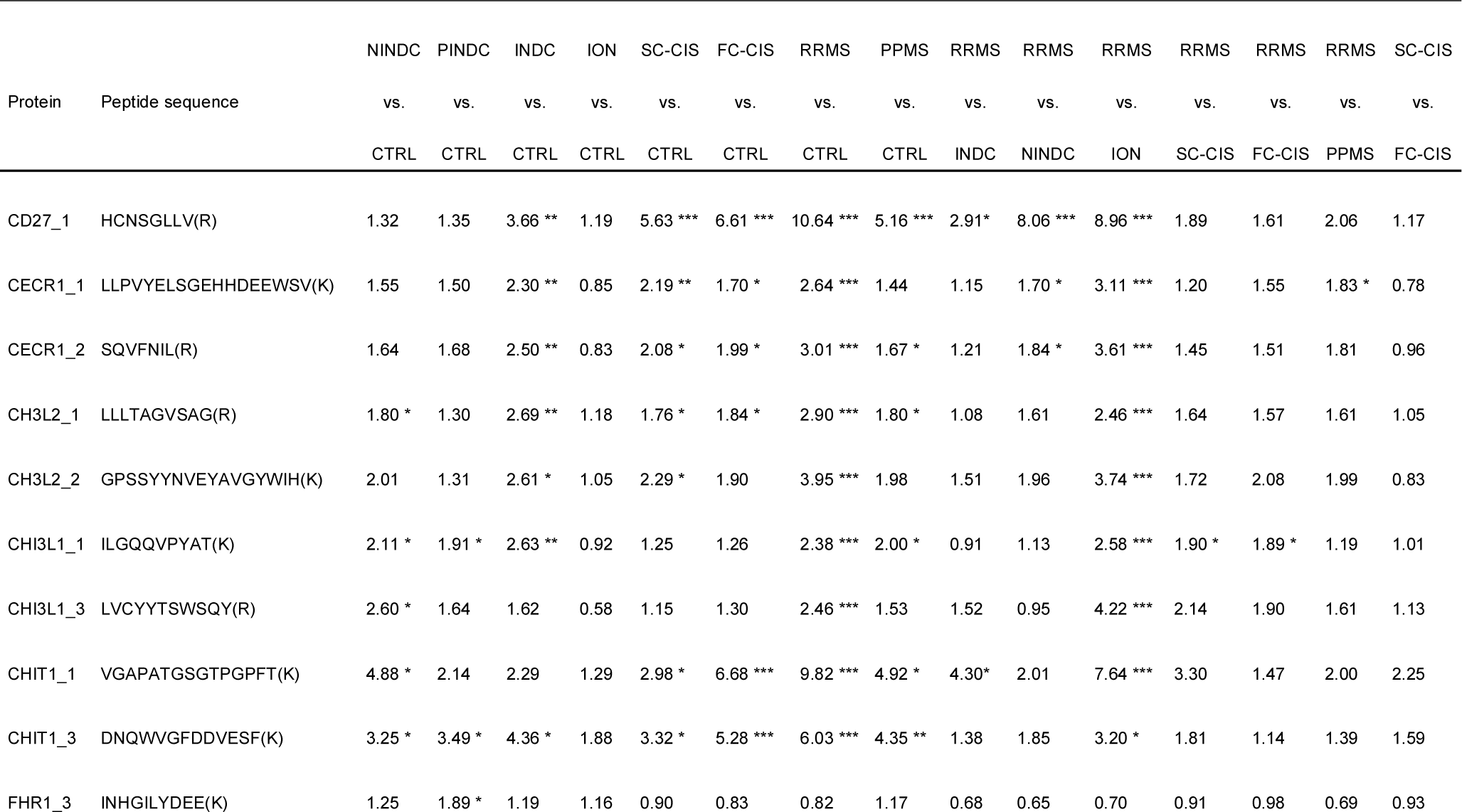
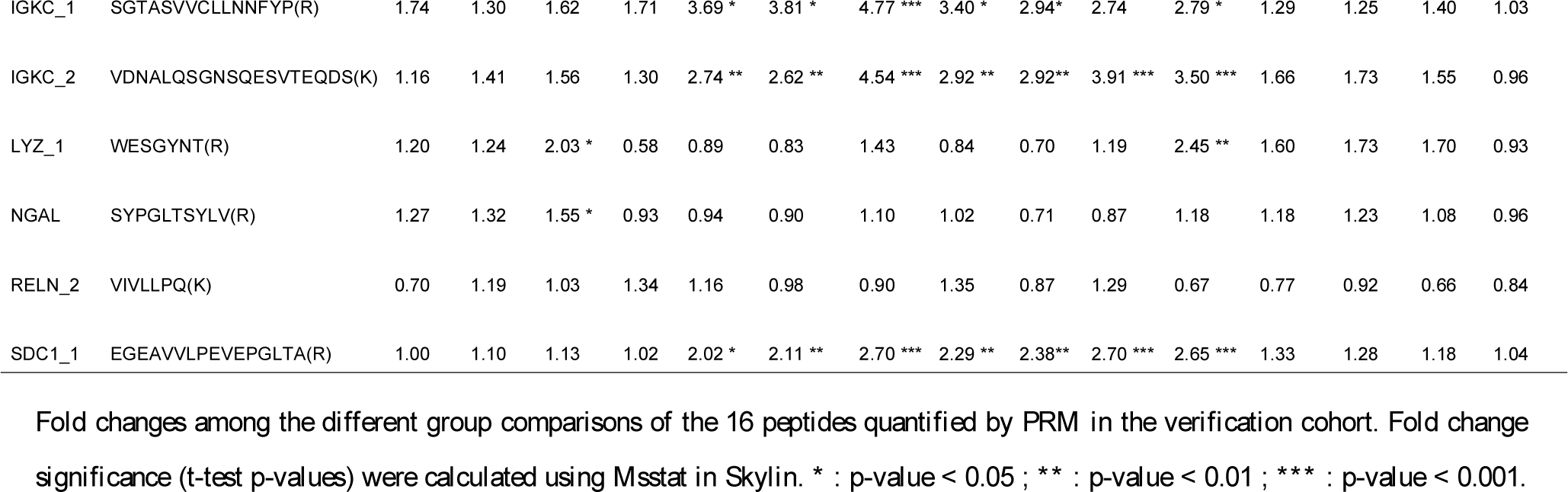
Value assigned to each statistical test and the global score for the proteins or peptides.

#### Targeted quantitative proteomics

For PRM analyses, CSF samples were immunodepleted of the 20 most abundant plasma proteins using the ProteoPrep 20 plasma protein immunodepletion kit, by performing a single immunodepletion cycle. Samples were then processed as described for label-free quantitative proteomics studies. For each protein, 1-3 peptides were selected for PRM analysis (Supplementary Table 4) based on the following criteria: proteotypic peptides of 7-25 amino acid length and carrying two or three charges that were identified and quantified in label-free analysis, absence of missed cleavage, methionine and proline and, as far as possible, with retention times providing a homogenous distribution along the chromatographic gradient. Heavy isotope-labelled versions of each monitored peptide (PEPotec SRM Grade 3, ThermoFisher Scientific) were spiked in the digested CSF samples at optimal dilution to obtain for each peptide a signal similar to those of the corresponding endogenous peptides. The determination of the optimal dilutions was performed by spiking a mixture of all heavy peptides at increasing dilutions (40X to 960,000X) in a CSF pool from patients with different neurological diseases. PRM analyses were performed using the Q-Exactive+ instrument used for data-dependent analyses (DDA). Each sample was analysed using the same LC gradient excepted 0-26 % B in 80 min followed by 26-52% B in 12 min and two different methods. For abundant peptides, data were acquired continuously throughout the nano-LC separation according to peptide retention time window determined in a first round of analysis and orbitrap resolution of 17,500 (at m/z 200), with isolation window of 2, target AGC value of 5^5^, maximum filling time of 100 ms and normalized collision energy of 26. For low abundant peptides, data were acquired continuously throughout the nano-LC separation according to peptide retention time window determined in a first round of analysis, orbitrap resolution of 35,000 (at m/z 200), isolation window of 1.5, target AGC value of 16, maximum filling time of 250 ms and normalized collision energy of 26.

For the second PRM assay (verification step), high-purity heavy isotope-labelled peptides (AQUA Ultimate, Thermofisher Scientific) from proteins that exhibited significant abundance difference between the groups in the first PRM assay were spiked in the digested CSF samples at optimal dilutions. Samples were analysed with short LC gradient (same buffer, 0-26 % B in 11 min followed by 26-52% B in 6 min). Data were acquired continuously throughout the nano-LC separation according to peptide retention time window determined in a first PRM analysis, with orbitrap resolution of 35,000 (at m/z 200), isolation window of 1.5, target AGC value of 1^5^, maximum filling times of 200 ms and normalized collision energy of 26. PRM data were analysed using Skyline (v. 3.6.0) (27).

### Primary cultures of rat oligodendrocyte precursor cells (OPCs)

Animals were handled according to protocols approved by the University of Montpellier ethics committee for animal use (CEEA LR 34, #7251). Brain cortices from WT Wistar rats at post-natal day 1 were dissected in Hank’s buffer (Gibco), supplemented with 0.01 M HEPES, 0.75% sodium bicarbonate (Gibco) and 1% penicillin/streptomycin. Cortices were enzymatically dissociated with papain (30 mg/mL in DMEM supplemented with 0.24 mg/mL N-acetylcysteine and 40 mg/mL DNase I) for 30 min at 37°C and then mechanically dissociated using a fire-narrowed Pasteur pipette. Dissociated cells were filtered on 40-mm filters to remove cell debris and plated in 75 mL flasks in DMEM supplemented with 10% foetal calf serum (FCS). After 12 days in culture, OPCs were harvested by overnight shaking (250 rpm, 37°C). After removal of contaminating microglial cells by differential adhesion to uncoated 100-mm culture dishes for 15 min, the cell suspension enriched in OPCs was centrifuged at 200 × g for 5 min and cells were resuspended in the same medium and plated on 100-mm culture dishes coated with 1.5 μg/ml poly-L-ornithine (Sigma). Two hours after seeding, the medium was replaced with modified Bottenstein Sato medium containing DMEM deprived of arginine (Arg) and lysine (Lys), 0.5% FCS, 2 mM L-glutamine, 5 μg/ml insulin, 30 nM sodium selenite, 100 μg/mL transferrin, 0.28 mg/mL albumin, 20 nM progesterone, 100 μM putrescine, 40 ng/ml triiodothyronine and 30 ng/mL L-thyroxine and supplemented with either heavy (H) amino-acids (L-[^13^C_6_-^15^N_4_]arginine (Arg10) and L-[^13^C_6_-^15^N_2_]lysine (Lys8) or mid-heavy (M) amino-acids (L-[^13^C_6_]arginine (Arg6) and L-[^2^H_4_]lysine (Lys4)), Euriso-top, Saint-Aubin, France). After 4 days, cultures were found to contain 85% of OPCs, as assessed by NG2 immunostaining.

### Cell treatment and media conditioning

Cells were washed five times with modified Bottenstein Sato medium without heavy amino acids and FCS and exposed to either vehicle or TNFα (10 ng/ml) or soluble Fas ligand (sFasL) in the same medium for 24 h. After the 24-h secretion period, conditioned media were collected, centrifuged at 200 × g for 5 min and then at 20,000 × g for 25 min to remove non-adherent cells and cell debris, respectively.

### Quantitative OPC secretome analysis

Proteins from OPC supernatants were precipitated using 10 % trichloroacetic acid on ice for 30 min. Precipitated proteins were spun down at 10,000 × g for 20 min and washed three times with diethyl ether to remove any remaining salt from the protein pellets. Precipitated proteins were resuspended in SDS sample buffer (62.5 mM Tris-HCl, pH 6.8, 2% SDS, 10% glycerol, 1% 2-mercaptoethanol and 0.005% bromophenol blue), separated on 12 % polyacrylamide gels and stained with PageBlue Protein Staining Solution (Fermentas). Gel lanes were cut into 15 equal gel pieces. After reduction (with 10 mM dithiothreitol at 56°C for 15 min) and alkylation (55 mM iodoacetamide at room temperature for 30 min), proteins were digested *in-gel* using trypsin (600 ng/band, Gold, Promega), as previously described [29]. The resulting peptides were analyzed online by nano-flow HPLC-nanoelectrospray ionization as previously described for CSF sample analysis. Raw data were analysed using the MaxQuant software (version 1.4.1.2) (25) and the Andromeda search engine [http://coxdocs.org/doku.php?id=maxquant:andromeda:start] against the complete rat proteome dataset (Uniprot KB). Relative protein quantifications in samples to be compared were performed based on the median SILAC ratios of at least two peptides, using MaxQuant with standard settings. Significance thresholds were calculated by using Perseus (www.maxquant.org) based on significance B with a p value of 0.01 for normalized peptide ratios.

### Measurement of OPC apoptosis

OPC cultures were fixed with 4% paraformaldehyde in PBS (for 10 min at 4 °C). Nuclei were then stained with 1 μg/mL Hoechst 33258 (Sigma) at room temperature for 10 min. Cells were then washed with PBS and distilled water and mounted in Mowiol under coverslips. Nuclear DNA staining was analysed by fluorescence imaging microscopy using an Axiophot2 microscope (Carl Zeiss, Le Pecq, France) equipped with epifluorescence. Apoptosis was estimated by counting the number of condensed or fragmented nuclei relative to the total number of nuclei (stained with Hoechst 33258) in at least nine different fields (about 350 cells per field) from three independent cultures.

### ELISA

CHI3L1 concentration was determined in CSF samples from CTRLs and RRMS patients using the MicroVue YKL-40 EIA kit (Quidel Corporation, San Diego, CA) after x4 dilution, otherwise according to the manufacturer instructions.

SDC1 and CD27 concentrations were determined in undiluted CSF samples from CTRL, RRMS and PPMS patients using the R-PLEX Human Syndecan-1 Assay and the and U-PLEX Human CD27 Assay, respectively (Meso Scale Discovery, Rockville, MD, USA) with a MESO QuickPlex SQ 120MM according to the manufacturer instructions. The CSF SDC1 and CD27 lower limit of detection (LLOD) were 4.2 pg/mL and 0.3 pg/mL, respectively.

### Immunocytochemistry

OPC-enriched cultures grown on glass coverslips were fixed at 6 days *in vitro* (DIV) with 2% paraformaldehyde for 15 min at 37°C, then rinsed three times for 10 min with PBS supplemented with BSA (0.5%) and glycine (0.1 M). They were permeabilized with triton X-100 (0.05%) for 5 min followed by 2 washes in PBS supplemented with 0.05% BSA (PBS-BSA). They were then incubated in a blocking solution (PBS-BSA containing 1% goat serum) for 1 h, then with the primary antibodies in the same solution overnight at 4°C (SDC1, Rabbit Thermofisher MA5-32600, 1:500 dilution; GFAP, mouse Sigma G3893, 1:500 dilution; O4, mouse R&D MAB1326, 1:1,000 dilution; MBP, mouse Ozyme BLE 808401, 1:1,000 dilution). After three washes, the cells were incubated with secondary antibodies at 1:1,000 dilution in PBS-BSA for 1 h at RT. Cells were then washed thrice in PBS-BSA, once in PBS and incubated for 10 min in PBS with Hoechst 33342 (2 μM, Thermo Scientific, Ref 62249). Coverslips were mounted on a slide in a mounting solution (Dako, Ref S3023) and immunofluorence images were captured with a Zess AxioImager Z1 Microscope equipped with an Apotome grid.

### Immunohistochemistry

For horseradish peroxidase (HRP) labelling, paraffin sections (4-µm tick) of fixed brain were subjected to antigen retrieval after quenching of endogenous peroxidase, by immersion in citrate/ EDTA buffer, pH 6 and heating (40 min at 100°C). Mouse anti-human SDC1 antibody (MI15, ThermoFischer) was used as primary antibody at 1:500 dilution. Biotinylated secondary antibody was raised against mouse IgGs. Immunoperoxidase reaction was performed using the avidin-biotin method and 3’,3’diaminobenzidine as chromogen and the ROCHE automatic immunostaining system (Benchmark ULTRA). Sections were then counterstained with Haematoxylin.

For double IHC staining, 3’,3’diaminobenzidine SDC1-labelled and unlabelled slices counterstained with Haematoxylin were further incubated for 1 h with the rabbit anti-human CHI3L1 antibody (Abcam, Ref ab77528, 1:500 dilution). After three washes, slices were incubated 30 min in AP One-Step Polymer anti-Mouse/Rabbit/Rat (Zytomed Systems, ref ZUC068) and labelled with Permanent AP Red (Zytomed Systems, ref ZUC001) according to the manufacturer’s instructions.

For immunofluorescence, fresh frozen brain slices on glass coverslips were incubated in PBS solution containing 10% heat inactivated goat serum (Vector Laboratories, Ref S-100) and 0.3% Triton X-100 for 20 min. They were then incubated overnight at 4°C in PBS containing 3% heat inactivated goat serum, 0.1% Triton X-100, and primary antibodies (SDC1 Rabbit Thermo Fisher Scientific, MA5-32600, 1:500 dilution; GFAP mouse Sigma G3893, 1:500 dilution). After extensive PBS washings, slices were incubated for 2 h with the Alexa Fluor^®^ 594-conjugated anti-rabbit antibody (Thermo Fisher Scientific, Ref A-11037, 1:1,000 dilution) and the Alexa Fluor^®^ 680-conjugated anti-mouse antibody (Thermo Fisher Scientific, Ref A-21057, 1:1,000 dilution) in PBS containing 3% heat inactivated goat serum, 0.1% Triton X-100 and Hoechst 33342 (2 μM, Thermo Fisher Scientific, Ref 62249). After three washes in PBS, glass coverslips were mounted on superfrost ultra plus slides (Thermo Fisher Scientific, Ref 10417002) using fluorescent mounting medium (Dako, Ref S3023). Immunofluorescence images were taken with an AxioImager Z1 microscope equipped with Apotome (Zeiss).

## Results

### Identification of candidate prognostic biomarkers of MS by quantitative proteomic analysis of CSF

We first compared the CSF proteome of 10 CTRLs, 10 SC-CIS, 10 FC-CIS and 10 RRMS patients matched in age (mean age 35.3 to 37.6 years), sex ratio (70% in all groups), and CSF protein level (mean CSF protein level 0.39 to 0.42 g/L, Table 1 and Fig. 1). As expected, SC-CIS, FC-CIS and RRMS samples showed a higher IgG index and the presence of OCBs in CSF, compared with CTRLs (Table 1). Samples from each patient were analysed in triplicate, yielding a total of 120 LC-MS runs. Quality of LC-MS data, assessed by a dispersion tree representing protein expression in each sample after missing value imputation and data normalization, showed a regular dispersion of the data together with proximity of technical replicates (Supplementary Fig. 2), thus indicating similar protein composition of all samples and reproducibility of analyses.

**Figure 1.**
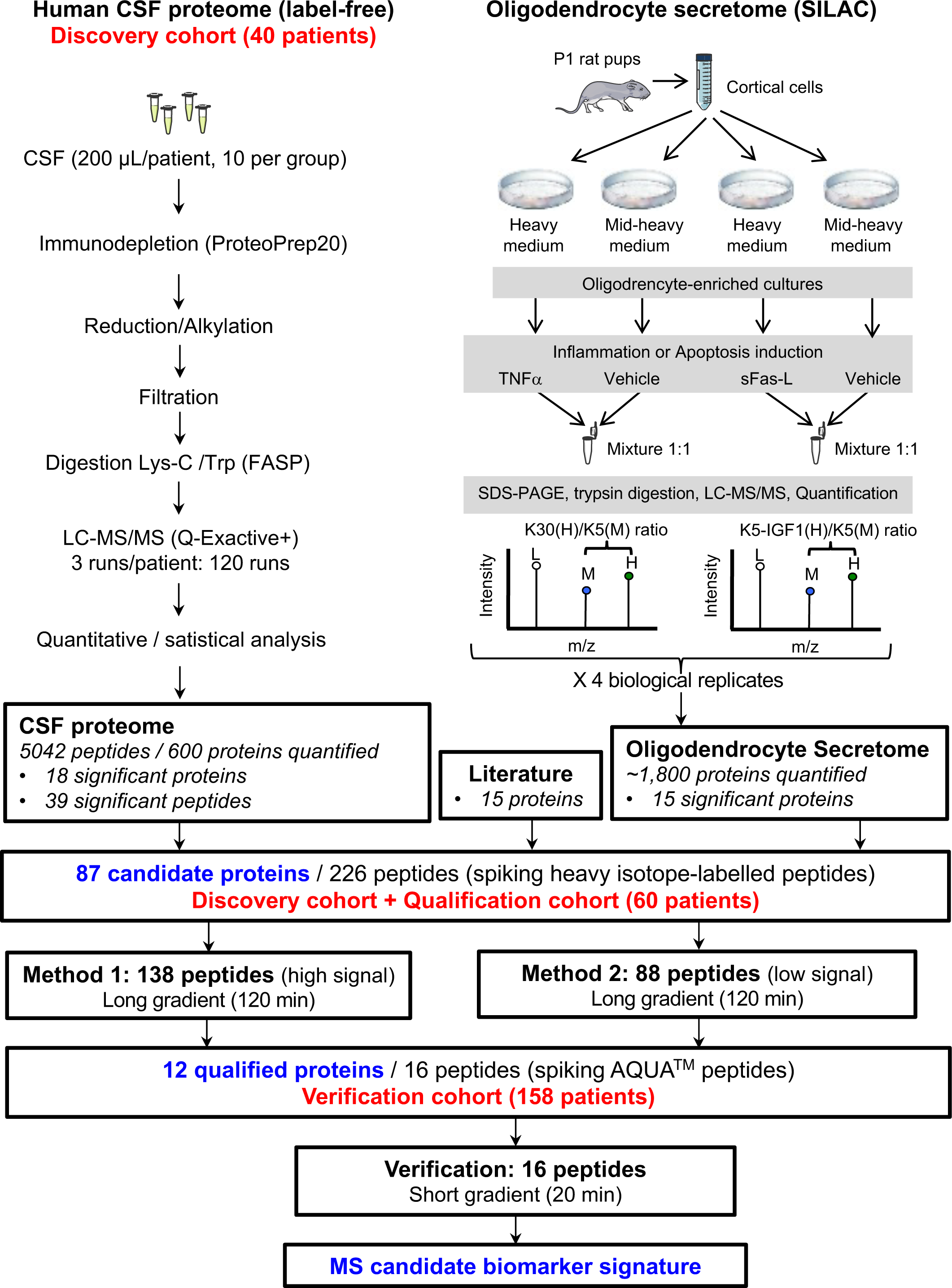
Schematic representation of the workflow used in the study.

Overall, a total of 5,042 unique peptides corresponding to 600 proteins were identified and quantified after data filtering. Comparing samples at protein level revealed 12 proteins exhibiting significant difference in abundance in RRMS *vs.* CTRL patients (Supplementary Table 1). These include 9 proteins more abundant (3 Ig kappa chains, CHI3L1, CHI3L2, chitotriosidase, adenosine deaminase CECR1 (Cat eye syndrome critical region protein 1), alpha-1-antichymotrypsin and protocadherin-17) and 3 less abundant (gamma-glutamyltransferase 7, desmocolin-1 and an Ig lambda chain) in RRMS samples, compared with CTRLs. Hierarchical clustering showed that these 12 proteins segregate both patient groups (Fig. 2A). Likewise, six proteins exhibited significant differences in abundance in CSF from FC-CIS *vs.* SC-CIS patients (Supplementary Table 1) and segregated both patient groups (Fig. 2B). Five of them (membrane frizzled-related protein, N-acetylgalactosamine-6-sulfatase, sodium/iodide cotransporter, alpha-N-acetylglucosaminidase and coagulation factor V) showed lower abundance in CSF from FC-CIS samples while brain acid soluble protein1 was overexpressed in CSF of FC-CIS patients. Analysis of data at peptide level revealed 39 additional proteins with at least one peptide exhibiting significantly different abundances in CSF from RRMS patients and CTRLs (Supplementary Table 2).

**Figure 2:**
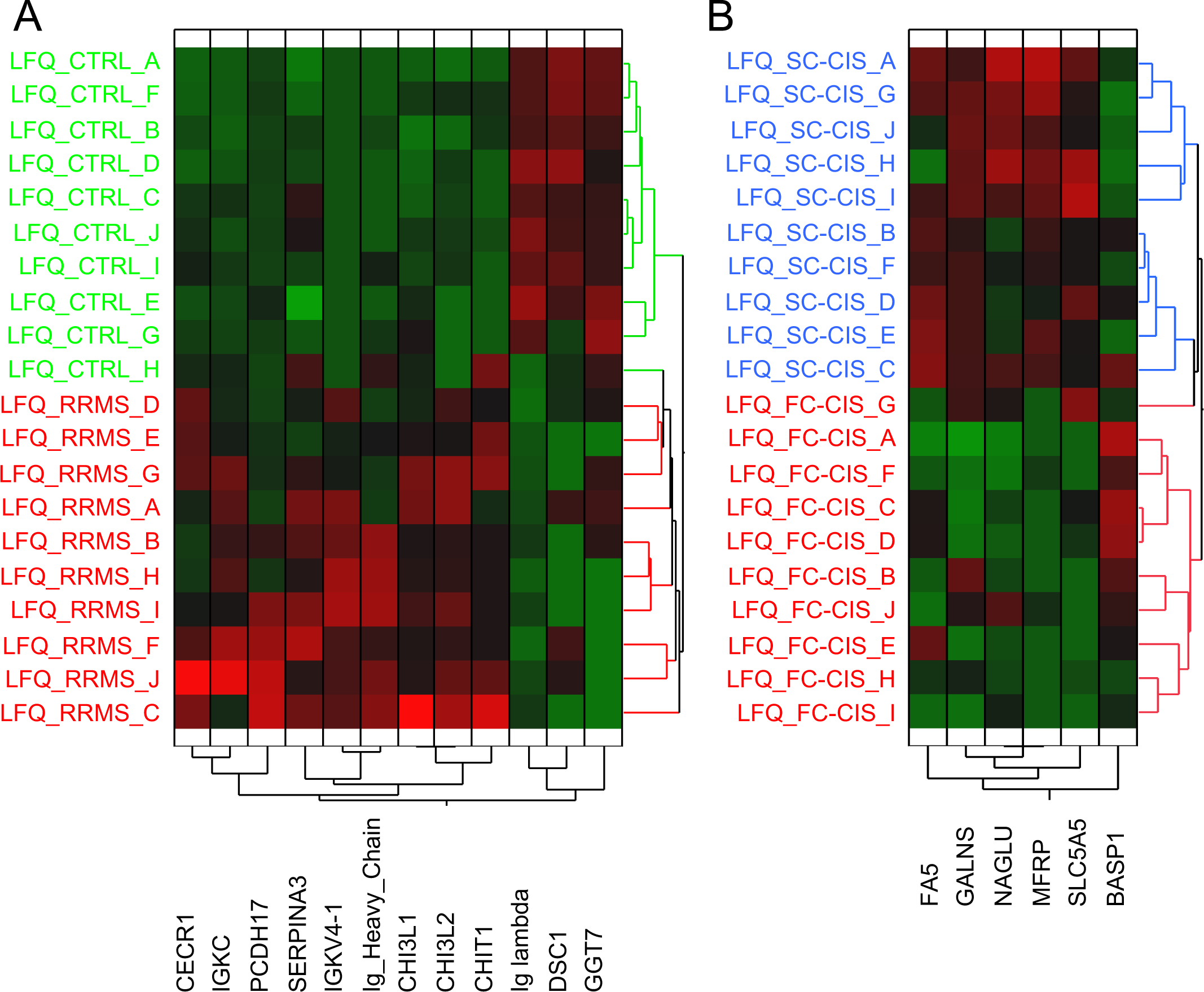
Results of label-free proteomic analysis of human CSF samples from the discovery cohort. Hierarchical clustering of the 12 proteins exhibiting difference in abundance in CSF from RRMS patients and CTRL (A) and of the 6 proteins exhibiting difference in abundance in CSF from FC-CIS and SC-CIS (B).

### Identification of potential biomarkers of inflammation and oligodendrocyte apoptosis by quantitative analysis of cultured OPC secretome

To identify additional candidate biomarkers reflecting two major pathological features of MS (inflammation and oligodendrocyte apoptosis), we next explored the modifications of the secretome of primary cultures enriched in rat OPCs elicited by exposing cultures with either TNFα or sFasL (both at 10 ng/mL) for 24 h, using the SILAC technology. As expected, exposure of OPCs to sFasL induced a significant increase in apoptotic OPCs in the cultures (46.5 ± 12.7 % condensed or fragmented nuclei in sFasL-treated cells cultures *vs.* 11.2± 3.0 % and 13.9 ± 5.6 % in vehicle-and TNFα-treated cultures, p<0.0001 and p<0.0001 (unpaired t-test), respectively, Supplementary Fig. 3).

We employed a double labelling procedure similar to those we previously used to analyse the secretome of primary cultured neurons (28) or astrocytes (29) to compare the relative abundance of proteins in conditioned media of OPCs treated with vehicle and either TNFα or sFasL, in order to avoid any bias in protein quantification related to uncomplete isotopic protein labelling. A total of 2,535 proteins were identified in OPC supernatant in four biological replicates comparing the secretome of vehicle and TNFα or sFasL-treated cells. Of these, 36 proteins showed significant difference in abundance (assessed by significance B) in the supernatant of OPCs treated with vehicle and TNFα in at least 2 out of 4 replicates and 19 proteins significant difference in abundance in the supernatant of vehicle and sFasL-treated OPCs (Supplementary Table 3). Fifteen human orthologs of differentially OPC-secreted proteins were identified in our quantitative proteomic analysis of human CSF samples (Supplementary Table 3).

### Qualification of candidate biomarkers by targeted quantitative proteomics

We next combined proteins showing difference in abundance at protein (18 proteins) or peptide level (39 proteins) in label-free quantitative analysis of patient CSF samples with proteins showing different abundance in OPC secretome upon exposure to TNFα or sFasL and identified in our proteomic analysis of human CSF (15 proteins) and 15 additional proteins previously identified as candidate biomarkers of MS (Supplementary Table 4, Figure 1) in a list of 87 proteins that were further analysed by PRM in the initial cohort and a new cohort of 60 patients (qualification cohort, Table 1) comprising CTRL, SC-CIS, FC-CIS, RRMS, PPMS and INDC (10 samples per group). For each of these proteins, we selected a maximum of three proteotypic peptides providing a good signal in our label-free analyses of human CSF, yielding a list of 226 peptides that were analysed by PRM (Supplementary Table 4, Figure 1). To improve the detection and relative quantification of peptides, digested CSF samples were spiked a mixture of heavy-isotope-labelled versions of these 226 peptides at concentrations yielding signals of similar intensities to those of the endogenous peptides. We first compared label-free and PRM RRMS/CTRL ratios for the 76 proteins quantified with both approaches in the discovery cohort. As shown on Supplementary Fig. 4A, a strong correlation of label-free and PRM RRMS/CTRL ratios was found in this cohort (Pearson coefficient, 0.86), thus validating our PRM strategy. Comparison RRMS/ CTRL PRM ratios in the discovery and qualification cohorts also indicated a good correlation (Pearson coefficient, 0.83, Supplementary Fig. 4B). Proteins selected for PRM analysis include the previously described MS biomarker CHI3L1 (4,5,30,31). Corroborating previous findings, CHI3L1 showed significant difference in abundance in CSF of CTRL and RRMS patients both in label-free shotgun analysis (discovery cohort, Supplementary Table 1 and 2) and PRM analysis (qualification cohort, Supplementary Table 5). Furthermore, CSF CHI3L1 concentration, determined by ELISA, was correlated with label-free and PRM CHI3L1 quantification in the discovery cohort (Pearson coefficients: 0.68 and 0.59, respectively, Supplementary Fig. 4C and 4D), further validating quantitative proteomics approaches used for MS biomarker discovery and verification. Out of the 226 peptides analysed, 16 peptides corresponding to 11 different proteins exhibited significant PRM ratios in RRMS *vs.* CTRLs, RRMS *vs*. PPMS or RRMS *vs.* INDCs comparisons (Table 2). These proteins include previously identified candidate biomarkers of MS such as CHI3L1, CHI3L2, chitotriosidase, IGKC and CD27 and novel candidate biomarkers of the disease such as the adenosine deaminase CECR1 and the proteoglycan syndecan-1 (SDC1, also known as plasma cell surface marker CD138, Table 2). None of them showed significant difference in abundance FC-CIS *vs*. SC-CIS patient (Table 2).

### Verification of qualified biomarkers by targeted quantitative proteomics

These 11 proteins were next quantified by PRM in a new cohort of 158 patients (verification cohort), including 30 CTRL, 13 NINDC, 13 PINDC, 13 INDC, 15 ION, 15 SC-CIS, 15 FC-CIS, 30 RRMS and 14 PPMS patients (Table 2). For this new PRM analysis, heavy isotope-labelled and high-purity (AQUA^TM^) versions of the 16 analysed peptides were spiked in the digested CSF samples for absolute quantification and determination of LOD and LOQ. Among the 11 proteins investigated in this second PRM analysis, eight exhibited differences in abundance between RRMS and CTRL or ION patients, or between patients with MS at any disease stage and other inflammatory and non-inflammatory neurological diseases. These include the previously identified MS biomarker CHI3L1, which showed an increased level not only in MS patients (at all disease stages) but also in INDC, NINDC and PINDC patients, when compared with CTRL and ION patients (Fig. 3). CHI3L2, CHIT1 and CECR1 showed similar differences in abundances between the different groups. Most importantly, CSF SDC1 and IGKC levels were more elevated in MS patients at all disease stages, compared with CTRL, INDC, NINDC, PINDC and ION patients, whereas neutrophil-associated gelatinase (NGAL) was more abundant in CSF from INDC, NINDC and PINDC patients compared to CIS and MS patients, suggesting that these proteins could distinguish between MS at any disease stage and other diseases (Fig. 3). CD27 concentration was likewise more elevated in CIS and MS patients but, contrasting with SDC1 and IGKC, it was also increased in INDC patients.

**Figure 3:**
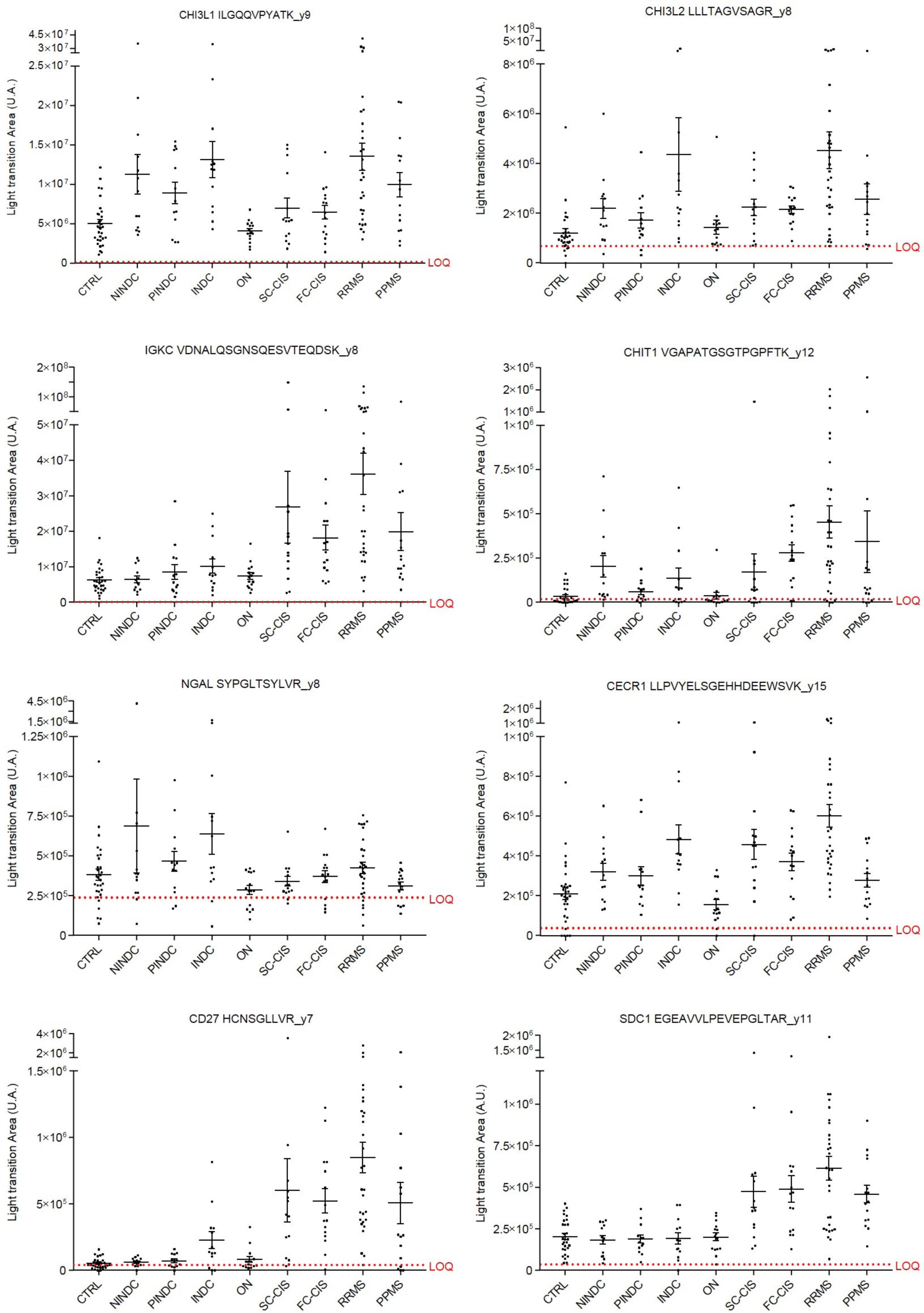
PRM analysis of peptides from eight candidate protein biomarkers in the verification cohort. Intensity (light transition area in arbitrary units (A.U.)) of eight peptides showing difference in abundance in CSF samples of the verification cohort is shown. The LOQ (limit of quantification) is indicated (red dotted line) for each peptide. Statistical analyses of group comparisons are provided in Table 3.

**Table 3:**
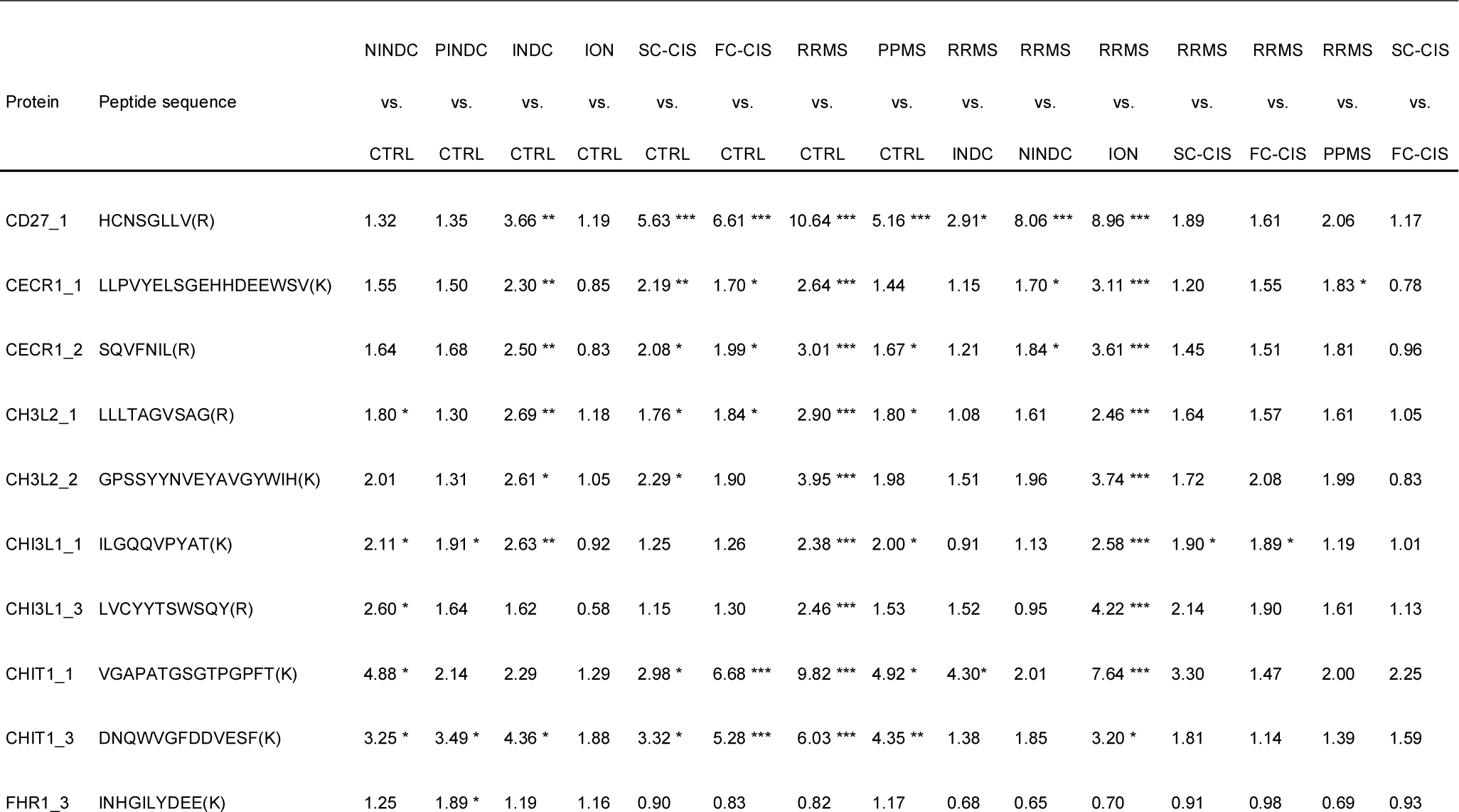
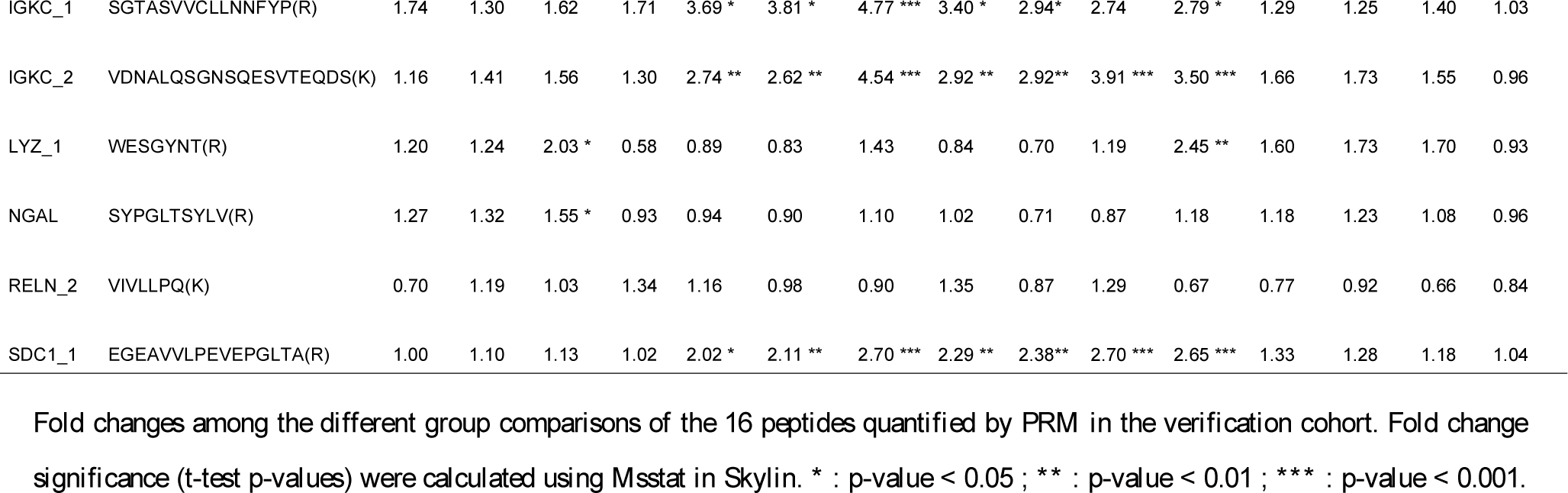
Peptide ratios measured by PRM in the verification cohort.

### Diagnostic value of verified candidate biomarkers

To further explore the diagnostic potential of these biomarkers, we investigated their sensitivity and specificity to discriminate MS patients from CTRLs and other neurological conditions. Receiver operating characteristic (ROC) curves showed that CHI3L1, CHI3L2, CHIT1, SDC1, IGKC, CD27 and CECR1 discriminate CTRL from RRMS patients as well as CTRL and ION patients from any neurological disease (SC-CIS, FC-CIS, RRMS, PPMS, INDC, PINDC or NINDC; Supplementary Table 6). Of these, CD27 has higher sensitivity and specificity in discriminating CTRL and RRMS (AUC=0.98, Fig. 4A and Supplementary Table 6), while the combination of CHI3L1, CHIT1 and SDC1 has higher sensitivity and specificity than each protein taken individually (AUC=0.88, Fig. 4B and Supplementary Table 6) in discriminating CTRL and ION from MS (at all disease stages), INDC, PINDC or NINDC, as assessed by multivariate analysis. On the other hand, CD27, SDC1, IGKC, CHI3L2, CHIT1 and CECR1 discriminate inflammatory CNS diseases (SC-CIS, FC-CIS, RRMS, PPMS, INDC) from other neurological diseases (PINDC or NINDC, Supplementary Table 6). Multivariate analysis showed that the combination of CD27 and SDC1 has higher sensitivity and specificity than each protein taken individually (AUC=0.91, Fig. 4C and Supplementary Table 6). Furthermore, SDC1 (AUC=0.85) is more efficient than IGKC (AUC=0.75), CD27 (AUC=0.76) and NGAL (AUC=0.69) to discriminate MS from INDC, and multivariate analysis showed that no combination of these markers is better than SDC1 alone to discriminate these groups (Fig. 4D and Supplementary Table 6). Finally, CECR1 (AUC=0.78) better differentiates RRMS from PPMS patients than NGAL (AUC=0.64), and multivariate analysis showed that the combination of these two markers is not better than CECR1 alone to discriminate these groups (Fig. 4E and Supplementary Table 6).

**Figure 4:**
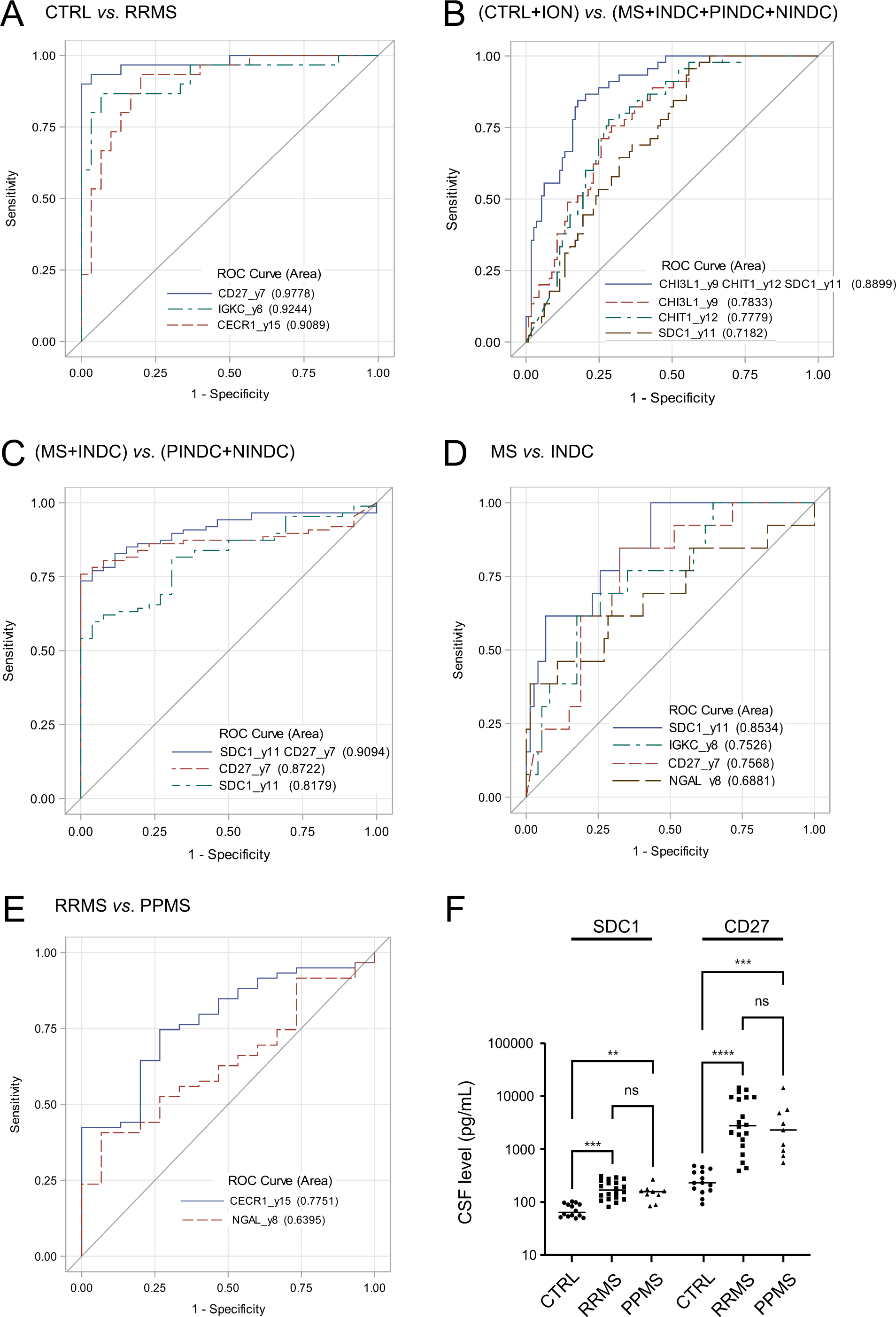
Diagnostic value of candidate MS biomarkers. A) ROC curves showing the diagnostic values of CD27 (AUC=0.98), IGKC (AUC=0.92) and CECR1 (AUC=0.91) to discriminate RRMS from CTRL. Multivariate analysis indicated that no combination has better diagnostic value than CD27 alone. B) ROC curves showing the diagnostic values of CHI3L1 (AUC=0.78), CHIT1 (AUC=0.78) and SDC1 (AUC=0.72) to discriminate CTRL and ION from MS, INDC, PINDC and NINDC. Multivariate analysis showed that only the combination of CHI3L1, CHIT1 and SDC1 has a better sensitivity and specificity to discriminate both groups (AUC=0.88). C) ROC curves showing the diagnostic value of CD27 (AUC=0.87) and SDC1 (AUC=0.82) to discriminate MS and INDC from PINDC and NINDC. Multivariate analysis showed that only the combination of CD27 and SDC1 has a better sensitivity and specificity to discriminate both groups (AUC=0.91) D) ROC curves showing the diagnostic values of SDC1 (AUC=0.85) and NGAL (AUC=0.69) to discriminate MS from INDC. Multivariate analysis indicated that no combination has better diagnostic value than SDC1 alone. E) ROC curves showing the diagnostic values of NGAL (AUC=0.68), CD27 (AUC=0.71), IGKC (AUC=0.70) and CECR1 (AUC=0.85) to discriminate RRMS from PPMS. Multivariate analysis indicated that no combination has better diagnostic value than CECR1 alone. F) CSF SDC1 and CD27 levels measured by ELISA in CTRL (n=14), RRMS (n=20) and PPMS (n=9) patients. **: p-value < 0.01; ***: p-value < 0.001; ****: p-value < 0.0001.

To further validate the results of the PRM approach, we measured SDC1 and CD27 by ELISA, and showed that these two markers are significantly increased in the CSF from RRMS and PPMS patients, as compared to CTRLs (Fig. 4F).

### Enhanced expression of SDC1 in MS brain

Given the potential of SDC1 to discriminate MS from other CNS inflammatory and non-inflammatory diseases, we explored its expression in brain slices from a CTRL and an RRMS patient by immunohistochemistry. Whereas SDC1 immunolabelling was not detected in CTRL brain, a strong SDC1 immunostaining was predominantly observed in round-shaped cells of the perivascular spaces (Fig. 5A). A weaker SDC1 staining was also found in cells of the parenchyma, especially in the vicinity of inflamed perivascular spaces. Furthermore, SDC1 staining was not colocalized with CHI3L1 immunostaining (Fig. 5A) observed in reactive astrocytes of MS brain (4). Corroborating these observations, immunofluorescence staining showed that SDC1 is not colocalized with GFAP in RRMS brain (Fig. 5B). Likewise, SDC1 was not detected in GFAP-positive astrocytes in rat primary cultures containing OPCs and astrocytes (Fig. 5C). In contrast, a strong SDC1 labelling was found in OPCs (O4+) and, to a lesser extent, in mature (MBP+) oligodendrocytes (Fig. 5C).

**Figure 5:**
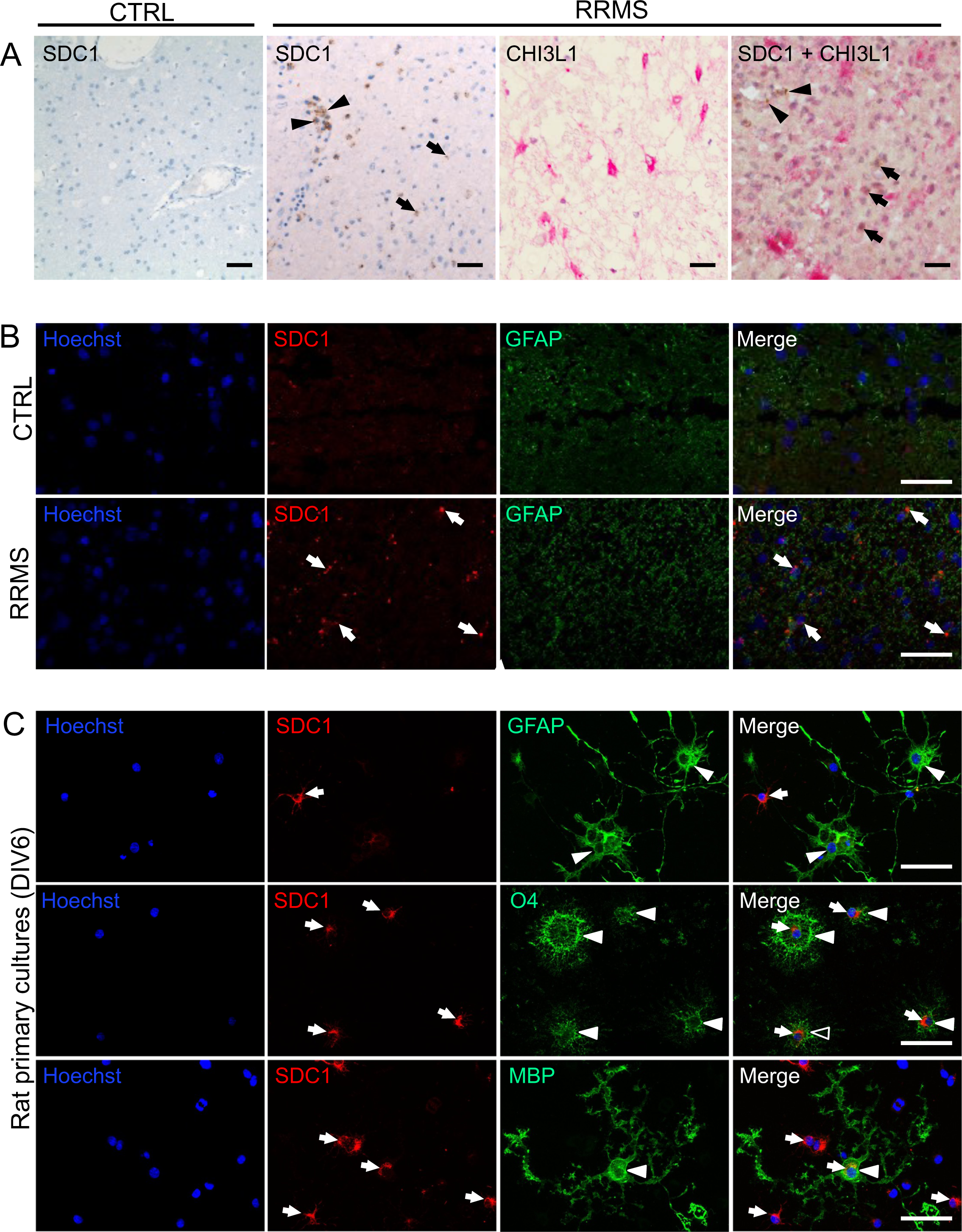
Expression of SDC1 in control and RRMS human brain and rat primary cultures. A) Immunohistochemistry of brain tissue showing predominant expression of SDC1 in cells located in the perivascular spaces (arrowheads) and a sparse expression in brain parenchyma (arrows) of RRMS brain but not in CTRL brain. CHI3L1 is strongly expressed in astrocytes (pink) from RRMS brain but shows no colocalization with SDC1 (brown) Scale bar: 100 µm. B) Immunofluorescence labelling showing a high expression of GFAP in activated astrocytes of RRMS brain as compared to CTRL brain. There is no colocalization of SDC1 (arrows) and GFAP in RRMS brain. Scale bar: 100 µm. C) Immunostaining of SDC1 (arrows) in rat primary cultured OPCs and astrocytes at 6 DIV showing a stronger expression of SDC1 in immature (O4+) than in mature (MBP+) oligodendrocytes, and its absence in astrocytes (GFAP+) (arrowheads). Scale bar: 100 µm.

## Discussion

Using two complementary proteomic strategies comparing i) the CSF proteomes of CTRL and RRMS patients and of CIS patients showing slow or fast conversion to RRMS and ii) the secretome of OPC cultures exposed or not to a pro-apoptotic or a pro-inflammatory treatment, we identified 72 candidate biomarkers of MS. They were combined with 15 previously described candidate biomarkers to generate a list of 87 proteins for further quantification by PRM in a new cohort comprising MS patients at different disease stages and control neurological diseases. The reliability of relative protein quantification by the label-free approach and PRM was supported by the correlation of the data obtained with both quantitative proteomics methods in the discovery cohort as well as the correlation between RRMS/CTRL PRM ratios in the discovery and qualification cohorts. Likewise, levels of the previously characterized MS biomarker (CHI3L1) measured by label-free approach, PRM and ELISA were strongly correlated, further supporting the relevance of our global and targeted quantitative proteomics assays used to identify and verify candidate biomarkers of MS.

PRM analysis of 226 peptides from the 87 candidate biomarkers qualified 11 of them for a second PRM round in a larger cohort of 158 patients that revealed a signature of eight biomarkers of potential interest in clinical practice. These include five previously described biomarkers of MS, such as CHI3L1, CHI3L2 and CHIT1 (31,32), IGKC, the constant region of the kappa free light chain of immunoglobulins, *i.e*. kFLC (17) and CD27 (33,34). All exhibit increased level in CSF of MS patients, consistent with previous findings. The three other proteins that passed the verification step include NGAL (also called lipocalin-2), an iron-binding protein involved in innate immunity and known to inhibit remyelination *in vitro* (35), SDC1, a cell surface proteoglycan bearing heparan and chondroitin sulphates that links the cytoskeleton to the extracellular matrix (36), and the secreted adenosine deaminase CECR1 that binds to cell surface proteoglycans and may play a role in the regulation of cell proliferation and differentiation independently of its adenosine deaminase activity (37). NGAL was identified as a candidate biomarker of MS in the analysis of OPC secretome but not in our label-free CSF proteome analyses, underpinning the complementarities of both approaches. Further, a decreased level of NGAL was measured in CSF of MS patients, compared to CTRLs, consistent with previous findings (38), while CSF SDC1 and CECR1 concentrations were found to be increased in MS in the label-free quantitative proteomic study (discovery step) and the two PRM verification steps. Further supporting the reliability of our results, increased levels of SDC1 and CECR1 were found in CSF of MS *vs.* CTRL patients in a previous shotgun CSF proteomic analysis (39), but no validation of these proteins as biomarkers of MS had so far been reported. Finally, none of the candidate biomarkers resulting from literature analysis (not identified in our initial proteomic screens) passed the PRM verifications step, underscoring the importance of biomarker candidate validation in different patient cohorts. Likewise, none of the eight identified biomarkers arise from the comparison of the CSF proteome from SC-CIS and FC-CIS patients and can thus be considered as biomarkers of disease activity, which remains a challenging issue. Previous proteomic studies dedicated to MS biomarker discovery, such as the pioneer study of Comabella *et al.*, actually compared CIS patients with normal MRI and absence of OCBs or patients with ION, to CIS patients with 3 or 4 Barkhof criteria and presence of OCBs (now considered as RRMS) (3,5). Only a recent study identified homeobox protein Hox-B3 (HOXB3) as candidate biomarker of conversion in CIS, but the results need further validation (40).

The eight biomarkers exhibit different sensitivities and specificities for MS but also complementary properties potentially useful for differential diagnosis of MS. Multivariate analysis indicated that a subset of them (CHI3L1, CHIT1 and SDC1) discriminate CTRLs and IONs from any other neurological diseases, including NINDCs. Accordingly, low CSF levels of these biomarkers might indicate the absence of any CNS diseases in patients complaining about neurological symptoms.

The adenosine deaminase (ADA2) CECR1 is another candidate biomarker of potential interest. CSF CECR1 concentration shows a large increase in RRMS patients, in comparison with CTRL, ION and PPMS patients. Further, multivariate analysis indicated that CSF CECR1 concentration discriminates RRMS patients from PPMS patients, suggesting that it is associated with the active phase of MS. Together with adenosine deaminase 1 (ADA1), CECR1 (ADA2) plays a key role in regulating the level of adenosine. It is secreted by monocytes undergoing differentiation into macrophages or dendritic cells (54). In turn, CECR1 induces T cell-dependent differentiation of monocytes into M2 macrophages and stimulates their proliferation through its recruitment at the cell surface via proteoglycans and adenosine receptors. In addition, CECR1 promotes production of pro-angiogenic factors, inhibits Th17 differentiation and stimulates Treg differentiation (55). CECR1 blockade by 2-chlorodeoxyadenosine (cladribine), a synthetic chlorinated deoxyadenosine analogue, modulates the immune responses during inflammation in selected cell types and provides targeted and sustained reduction of circulating T and B lymphocytes. Interestingly, cladribine has been developed as a treatment for active RRMS, indicating that CECR1 is not only a biomarker of the active phase of MS but also a therapeutic target (56,57).

Comparing CNS inflammatory diseases, including MS at all stages, with NINDCs and PINDCs showed that CD27, in combination with SDC1 discriminates both groups, indicating that this biomarker combination can be considered as biomarker of central inflammation. CD27 was also the most accurate biomarker discriminating RRMS from CTRLs. Interestingly, MS is characterized by the accumulation of B cells in the CSF. Most of them are memory B cells, with a high expression of CD27 at their surface, or short-lived plasma blasts expressing CD138 (SDC1) and CD38 (41). This specific CSF profile and the strong expression of SDC1 at the surface of plasmocytes and plasma blasts present in ectopic lymphoid follicles in the meninges could contribute, at least in part, to the increased concentration of SDC1 in the CSF (42). This could partially explain the results of our multivariate analysis revealing SDC1 as the only biomarker discriminating MS at all disease stages from other CNS inflammatory conditions (INDC). Although a recent study comparing CSF SDC1 level by ELISA in Neuromyelitis optica (NMO), MS and CTRLs revealed an increase in SDC1 concentration in NMO, but not in MS (43), these observations resulted from the analysis of a small group of patients (n = 12) based on a low sensitivity of the ELISA, compared with our PRM approach and with the high-sensitivity ELISA used in our study.

In the CNS, SDC1 is also highly expressed by choroidal epithelial cells, where it reduces leukocyte recruitment to the brain across the choroid plexuses (44). In the myelin oligodendrocyte glycoprotein–induced experimental autoimmune encephalomyelitis (MOG-EAE) MS model, SDC1 knockout mice show enhanced disease severity and impaired recovery, suggesting a protective role of SDC1. SDC1 seems to play a key role in blood brain barrier (BBB) integrity, which is altered in inflammatory brain disorders, including MS (45). On the other hand, SDC1 has been identified as a receptor that binds to CHI3L1 through heparan sulphate residues of its extracellular part and mediates CHI3L1-operated signalling involved in inflammation and cancer (46,47). Likewise, SDC1 has been suggested as an endothelial cell receptor mediating CHI3L1-induced angiogenesis (48,49). Association of CHI3L1 to SDC1 promotes recruitment of integrin αvβ3 to the membrane of vascular endothelial cells (48) and of integrin αvβ5 recruitment in tumor cells (49), leading to engagement of FAK and ERK1/2 signalling and VEGF expression. CHI3L1 also increases the expression of MMP9, CCL2, and CXCL2 *via* SDC1 (50), an additional process potentially contributing to BBB leakage in inflammation. Collectively, these results suggest that SDC1 exerts both beneficial and deleterious influences on MS that depend, at least in part, on the binding of CHI3L1 or other growth factors and chemokines to its heparan sulphate chains (51). Further supporting the potential influence of the CHI3L1/SDC-operated pathways, SDC1 polymorphism has been associated with MS, specifically in women suffering from either PPMS or RRMS (52). In the present study, we found a predominant expression of SDC1 in OPCs and mature oligodendrocytes but not in astrocytes, suggesting that SDC1 acts as a receptor of CHI3L1 released by activated astrocytes. This is consistent with previous data from Bansal *et al.* showing that the expression of syndecan-1 and −3 is higher in the oligodendrocyte lineage cells than in astrocytes (53). Analysis of CHI3L1-SDC1 interaction and associated signalling in oligodendrocytes from MS brain certainly warrants further exploration to better understand their role in the pathophysiology of MS.

In conclusion, the present study using three different cohorts comprising CTRL, MS patients and patients with other inflammatory and non-inflammatory neurological diseases, provides one of the most comprehensive proteomic studies dedicated to MS biomarker discovery and validation currently available. It identified and validated CECR1, a therapeutic target of MS, as a marker of the active phase of the disease, and SDC1 as novel biomarker that allows differential diagnosis of MS versus other neurological disorders, including INDCs.

## Acknowledgments

This study was supported by grants from the Fondation pour l’Aide à la Recherche sur la Sclérose en Plaques (ARSEP Foundation), the University of Montpellier, the Centre National de la Recherche Scientifique, The Institut National de la Santé et de la Recherche Médicale and la Région Languedoc-Roussillon. Geoffrey Hinsinger was recipient of fellowships from ARSEP and the Hospital of Nimes. Lucile du Trieu de Terdonck was recipient of fellowships from ARSEP and la Region Languedoc-Roussillon. Mass spectrometry analyses were performed using the facilities of the Functional Proteomics Platform of Montpellier Languedoc-Roussillon.

