## Supplementary Material for "Syndecan-1 as specific cerebrospinal fluid biomarker of multiple sclerosis"

**Supplementary Figure 1: Schema illustrating the patient groups used in the study.**

The different patient groups used in the study, the course of the disease and of symptoms are illustrated. The red arrow indicates the disease stage in which the lumbar puncture was done. Conversion is defined by the presence of a relapse or new T2 or T1-gadolinium enhancing lesions. The red arrows show the time of lumbar puncture for cerebrospinal fluid collection. OCB: presence oligoclonal band in CSF; MRI: MRI criteria for patients in the different groups. CTRL: symptomatic controls; ION: isolated optic neuritis; PINDC: peripheral inflammatory neurological disease controls; NINDC: non-inflammatory neurological disease controls, INDC: inflammatory neurological disease controls; CIS: clinically isolated syndrome; FC-CIS: fast conversion to MS (<1 year) after a CIS; SC-CIS: slow conversion to MS (>2 years) after CIS; RRMS: relapsing remitting multiple sclerosis patients; PPMS: patients with progressive form of multiple sclerosis without period of symptoms remission.

**Supplementary Figure 2. Analysis of label-free proteomic data quality using a dispersion tree approach.**

Each sample of the discovery cohort was analyzed in triplicate by label-free quantitative shotgun proteomics. The protein intensities in each sample were represented by an expression vector and the Euclidian distance between vectors was calculated. The resulting distance matrix was used to perform a clustering of all the experiments and the data dispersion was analyzed. The proximity of branches in the dispersion tree representing protein expression in each sample after missing value imputation and data normalization indicates low variability of data and similar composition of samples. X in red: barycenter. X in blue: geometric center.

**Supplementary Figure 3. Apoptosis induced by TNFα or sFasL in rat primary cultures of OPCs.** Primary cultures rat OPCs were exposed to TNFα or sFasL (both at 10 ng/ml) for 24 h. Percentage of apoptotic nuclei (condensed or fragmented nuclei) were compared with the vehicle condition.

**Supplementary Figure 4. Correlation of label-free quantitative proteomics, targeted quantitative proteomics, and ELISA data.**

A) PRM RRMS/CTRL ratios (ratios of peak areas, assessed by using the Skyline software) of differentially expressed proteins (n=76) in the discovery cohort (10 RRMS and 10 CTRL) were compared to their ratios determined by label-free quantitative proteomics (LFQ intensities assessed by the MaxQuant software). B) Comparison of PRM RRMS/CTRL ratios in the discovery and verification cohorts containing each 10 RRMS and 10 CTRL. C and D) CHI3L1 concentration, assessed by ELISA, was compared to the relative intensity of label-free (C) and PRM (D) CHI3L1 signals in CSF samples of the discovery cohort.

**Supplementary Table 1: Proteins exhibiting significant difference in abundance in the CSF from RRMS vs. CTRL and FC-CIS vs. SC-CIS patients.**

| Accession number | Gene name | Protein name | Groups | Unique peptide | Fold change | Best Test | SAM | Limma | Nfold | AUC ROC | Global Score |
|---|---|---|---|---|---|---|---|---|---|---|---|
| P01625 | IGKV4-1 | Ig kappa chain V-IV region Len | RRMS vs. CTRL | 3 | 453 | 1 | 1 | 1 | 0.5 | 1 | 4.5 |
| P01766 | IGHV3-13 | Ig heavy chain V-III region BRO | RRMS vs. CTRL | 2 | 37 | 1 | 1 | 1 | 0.5 | 1 | 4.5 |
| Q15782-6 | CHI3L2 | Chitinase-3-like protein 2 | RRMS vs. CTRL | 10 | 5.58 | 1 | 1 | 1 | 0.5 | 1 | 4.5 |
| P01834 | IGKC | Ig kappa chain C region | RRMS vs. CTRL | 5 | 5.07 | 1 | 1 | 1 | 0.5 | 1 | 4.5 |
| B4E3Q4 | CECR1 | Adenosine deaminase CECR1 | RRMS vs. CTRL | 8 | 2.51 | 1 | 1 | 1 | 0.5 | 1 | 4.5 |
| P36222 | CHI3L1 | Chitinase-3-like protein 1 | RRMS vs. CTRL | 19 | 1.65 | 1 | 1 | 1 | 0.5 | 1 | 4.5 |
| Q9UJ14 | GGT7 | Gamma-glutamyltransferase 7 | RRMS vs. CTRL | 4 | 0.61 | 1 | 1 | 1 | 0.5 | 1 | 4.5 |
| P04208 | IGLV1-47 | Ig lambda chain V-I region WAH | RRMS vs. CTRL | 1 | 0.05 | 1 | 1 | 1 | 0.5 | 1 | 4.5 |
| Q08554-2 | DSC1 | Desmocollin-1 | RRMS vs. CTRL | 8 | 0.32 | 0.25 | 1 | 1 | 0.5 | 1 | 3.75 |
| Q9BY79-2 | MFRP | Membrane frizzled-related protein | FC-CIS vs. SC-CIS | 3 | 0.66 | 1 | 1 | 1 | 0.5 | 1 | 3.5 |
| Q13231-3 | CHIT1 | Chitotriosidase-1 | RRMS vs. CTRL | 14 | 5.51 | 1 | 0.25 | 0.25 | 0.5 | 1 | 3 |

| | | | | | | | | | | | |
|---|---|---|---|---|---|---|---|---|---|---|---|
| P34059 | GALNS | N-acetylgalactosamine-6-sulfatase | FC-CIS vs. SC-CIS | 4 | 0.53 | 0.25 | 1 | 0.25 | 0.5 | 1 | 2.75 |
| O14917-2 | PCDH17 | Protocadherin-17 | RRMS vs. CTRL | 8 | 2.8 | 0.25 | 0.25 | 0.25 | 0.5 | 1 | 2.25 |
| P01011 | SERPINA3 | Alpha-1-antichymotrypsin | RRMS vs. CTRL | 21 | 1.74 | 0.25 | 0.25 | 0.25 | 0.5 | 1 | 2.25 |
| Q92911 | SLC5A5 | Sodium/iodide cotransporter | FC-CIS vs. SC-CIS | 4 | 0.65 | 0.25 | 0.25 | 0.25 | 0.5 | 1 | 2 |
| P54802 | NAGLU | Alpha-N-acetylglucosaminidase | FC-CIS vs. SC-CIS | 16 | 0.63 | 0.25 | 0.25 | 0.25 | 0.5 | 1 | 2 |
| P12259 | F5 | Coagulation factor V | FC-CIS vs. SC-CIS | 20 | 0.66 | 0.25 | 0.25 | 0.25 | 0.5 | 0.5 | 1.5 |
| P80723 | BASP1 | Brain acid soluble protein 1 | FC-CIS vs. SC-CIS | 7 | 1.53 | 0.25 | 0.25 | 0 | 0.5 | 0.5 | 1.25 |

The results of quantification at the protein level using the LFQ algorithm of Maxquant software and of statistical analysis based on five tests are indicated. Scores from individual tests (0, 0.25, 0.5, 1) and global scores (0 to 4.5) were calculated as described in the Materials and Methods section. Fold changes between groups calculated as ratios of normalized intensities and the number of unique peptides identified by mass spectrometry are shown for each protein.

**Supplementary Table 2: Peptides significantly up or down regulated in CSF from RRMS *vs.* CTRL and FC-CIS *vs.* SC-CIS patients and selected for the first PRM analysis (qualification step).**

| Protein ID | Gene name | Protein name | Sequence | Best Test | Limma | SAM | Nfold | AUC | Global Score | Comparison | Nfold |
|---|---|---|---|---|---|---|---|---|---|---|---|
| B4E3Q4 | CECR1 | Adenosine deaminase CECR1 | SQVFNILR | 0.25 | 1 | 1 | 0.50 | 1.0 | 3.75 | CTRL / RRMS | 3.98 |
| B4E3Q4 | CECR1 | Adenosine deaminase CECR1 | TLIFPPSMHFFQAK | 1 | 1 | 1 | 0.50 | 1.0 | 4.50 | CTRL / RRMS | 2.85 |
| P01011 | SERPINA3 | Alpha-1-antichymotrypsin | LINDYVK | 0.25 | 1 | 1 | 0.50 | 0.5 | 3.25 | CTRL / RRMS | 3.31 |
| P02749 | APOH | Beta-2-glycoprotein 1 | WSPELPVCAPIICPPPSIPTFATLR | 0.25 | 0.25 | 0.25 | 0.50 | 1 | 2.25 | SC-CIS / FC-CIS | 1.69 |
| Q9NQ79-2 | CRTAC1 | Cartilage acidic protein 1 | NVASGEMNSVLEILYPR | 0.25 | 1 | 0.25 | 0.50 | 1.0 | 3.00 | CTRL / RRMS | 0.26 |
| P26842 | CD27 | CD27 antigen | HCNSGLLVR | 1 | 1 | 1 | 0.50 | 1.0 | 4.50 | CTRL / RRMS | 6.45 |
| Q8N3J6-2 | CADM2 | Cell adhesion molecule 2 | VDRSDDGVAVICR | 0.25 | 0.25 | 0.25 | 0.50 | 1 | 2.25 | SC-CIS / FC-CIS | 1.45 |
| Q9NTU7 | CBLN4 | Cerebellin-4 | PVISAFAGDK | 0.25 | 1 | 1 | 0.50 | 1.0 | 3.75 | CTRL / RRMS | 0.12 |
| Q9NTU7 | CBLN4 | Cerebellin-4 | PVISAFAGDK | 0.25 | 0.25 | 0.25 | 0.50 | 1 | 2.25 | SC-CIS / FC-CIS | 5.05 |
| P36222 | CHI3L1 | Chitinase-3-like protein 1 | EAGTLAYYEICDFLR | 1 | 1 | 1 | 0.50 | 1.0 | 4.50 | CTRL / RRMS | 1.65 |

| | | | | | | | | | | | |
|---|---|---|---|---|---|---|---|---|---|---|---|
| P36222 | CHI3L1 | Chitinase-3-like protein 1 | QLAGAMVWALDLDDFQGSFCGQDLR | 0.25 | 1 | 1 | 0.50 | 1.0 | 3.75 | CTRL / RRMS | 2.13 |
| P36222 | CHI3L1 | Chitinase-3-like protein 1 | QLLLSAALSAGK | 0.25 | 1 | 1 | 0.50 | 1.0 | 3.75 | CTRL / RRMS | 2.46 |
| P36222 | CHI3L1 | Chitinase-3-like protein 1 | THGFDGLDLAWLYPGR | 1 | 1 | 1 | 0.50 | 1.0 | 4.50 | CTRL / RRMS | 1.91 |
| Q15782-6 | CHI3L2 | Chitinase-3-like protein 2 | GPSSYYNVEYAVGYWIHK | 1 | 1 | 1 | 0.50 | 1.0 | 4.50 | CTRL / RRMS | 4.96 |
| Q15782-6 | CHI3L2 | Chitinase-3-like protein 2 | LLLTAGVSAGR | 1 | 1 | 1 | 0.50 | 1.0 | 4.50 | CTRL / RRMS | 5.42 |
| Q15782-6 | CHI3L2 | Chitinase-3-like protein 2 | LQDQQVPYAVK | 1 | 1 | 1 | 0.50 | 1.0 | 4.50 | CTRL / RRMS | 4.79 |
| Q15782-6 | CHI3L2 | Chitinase-3-like protein 2 | LVCYFTNWSQDR | 1 | 1 | 1 | 0.50 | 1.0 | 4.50 | CTRL / RRMS | 2.51 |
| Q15782-6 | CHI3L2 | Chitinase-3-like protein 2 | NHNFDGLDVSWIYPDQK | 0.25 | 1 | 1 | 0.50 | 1.0 | 3.75 | CTRL / RRMS | 4.61 |
| Q13231-3 | CHIT1 | Chitotriosidase-1 | DNQWVGFDDVESFK | 1 | 1 | 1 | 0.50 | 1.0 | 4.50 | CTRL / RRMS | 11.61 |
| P10645 | CHGA | Chromogranin-A | PSPMPVSQECFETLR | 0.25 | 0.25 | 0.25 | 0.50 | 1 | 2.25 | SC-CIS / FC-CIS | 2.71 |
| Q03591 | CFHR1 | Complement factor H-related protein 1 | STDTSCVNPPTVQNAHILSR | 0.25 | 0.25 | 0.25 | 0.50 | 1 | 2.25 | SC-CIS / FC-CIS | 2.41 |
| Q02246 | CNTN2 | Contactin-2 | WGCAAAGK | 0.25 | 1 | 0.25 | 0.50 | 1.0 | 3.00 | CTRL / RRMS | 0.21 |
| Q08554-2 | DSC1 | Desmocollin-1 | VTIFTVPENCR | 0.25 | 1 | 1 | 0.50 | 1.0 | 3.75 | CTRL / RRMS | 0.31 |
| Q08554-2 | DSC1 | Desmocollin-1 | VTIFTVPENCR | 0.25 | 0.25 | 0.25 | 0.50 | 1 | 2.25 | SC-CIS / FC-CIS | 0.61 |

| | | | | | | | | | | | |
|---|---|---|---|---|---|---|---|---|---|---|---|
| P15924 | DSP | Desmoplakin | LNDSILQATEQR | 0.25 | 1 | 1 | 0.50 | 1.0 | 3.75 | CTRL / RRMS | 0.33 |
| P15924 | DSP | Desmoplakin | QLQNIIQATSR | 0.25 | 1 | 1 | 0.50 | 1.0 | 3.75 | CTRL / RRMS | 0.35 |
| P14625 | HSP90B1 | Endoplasmin | FQSSHHPTDITSLDQYVER | 0.25 | 1 | 0.25 | 0.50 | 1.0 | 3.00 | CTRL / RRMS | 4.77 |
| P02671-2 | FGA | Fibrinogen alpha chain | GSESGIFTNTK | 0.25 | 0.25 | 0.25 | 0.50 | 1 | 2.25 | SC-CIS / FC-CIS | 0.21 |
| P15328 | FOLR1 | Folate receptor alpha | DVSYLYR | 0.25 | 0.25 | 0.25 | 0.50 | 1 | 2.25 | SC-CIS / FC-CIS | 0.37 |
| P47929 | LGALS7 | Galectin-7 | SSLPEGIRPGTVLR | 1 | 0.25 | 0.25 | 0.50 | 1.0 | 3.00 | CTRL / RRMS | 0.61 |
| O00451 | GFRA2 | GDNF family receptor alpha-2 | QTILPSCSYEDK | 0.25 | 0.25 | 0.25 | 0.50 | 1 | 2.25 | SC-CIS / FC-CIS | 2.54 |
| P48058-2 | GRIA4 | Glutamate receptor 4 | NTDQEYTAFR | 0.25 | 0.25 | 0.25 | 0.50 | 1 | 2.25 | SC-CIS / FC-CIS | 2.71 |
| F8W785 | GOLIM4 | Golgi integral membrane protein 4 | NNDVWQNHEAVPGR | 1 | 1 | 1 | 0.50 | 1.0 | 4.50 | CTRL / RRMS | 0.32 |
| P01834 | IGKC | Ig kappa chain C region | SGTASVVCLLNNFYPR | 0 | 1 | 1 | 0.50 | 1.0 | 3.50 | CTRL / RRMS | 4.74 |
| P01598 | | Ig kappa chain V-I region EU | DIQMTQSPSTLSASVGDR | 1 | 1 | 1 | 0.50 | 1.0 | 4.50 | CTRL / RRMS | 8.42 |
| P01620 | | Ig kappa chain V-III region HIC | LLIYGASSR | 1 | 1 | 1 | 0.50 | 1.0 | 4.50 | CTRL / RRMS | 3.77 |
| P06311 | | Ig kappa chain V-III region IARC/BL41 | ASQSVSSNLAWYQQK | 1 | 1 | 0.25 | 0.50 | 1.0 | 3.75 | CTRL / RRMS | 3.22 |
| P01625 | IGKV4-1 | Ig kappa chain V-IV region Len | DIVMTQSPDSLAVSLGER | 1 | 1 | 1 | 0.50 | 1.0 | 4.50 | CTRL / RRMS | 8.94 |

| | | | | | | | | | | | |
|---|---|---|---|---|---|---|---|---|---|---|---|
| P01701 | | Ig lambda chain V-I region NEW | RPSGIPDR | 1 | 1 | 1 | 0.50 | 1.0 | 4.50 | CTRL / RRMS | 3.24 |
| P04208 | | Ig lambda chain V-I region WAH | QSVLTQPPSASGTPGQR | 1 | 1 | 1 | 0.50 | 1.0 | 4.50 | CTRL / RRMS | 3.63 |
| P04208 | | Ig lambda chain V-I region WAH | SGTSASLAISGLR | 1 | 1 | 1 | 0.50 | 1.0 | 4.50 | CTRL / RRMS | 5.64 |
| P01714 | | Ig lambda chain V-III region SH | SELTQDPAVSVALGQTVR | 1 | 1 | 1 | 0.50 | 1.0 | 4.50 | CTRL / RRMS | 4.97 |
| B9A064 | IGLC2 | Ig lambda-2 chain C regions | ATLVCLISDFYPGAVTVAWK | 0.25 | 1 | 1 | 0.50 | 1.0 | 3.75 | CTRL / RRMS | 18.40 |
| B9A064 | IGLC2 | Ig lambda-2 chain C regions | SYSCQVTHEGSTVEK | 1 | 1 | 1 | 0.50 | 1.0 | 4.50 | CTRL / RRMS | 7.13 |
| Q9Y6R7 | FCGBP | IgGFc-binding protein | PAGWQVGGAQGCGECVSK | 0.25 | 0.25 | 0.25 | 0.50 | 1 | 2.25 | SC-CIS / FC-CIS | 0.60 |
| J3QQR8 | ICAM2 | Intercellular adhesion molecule 2 | SFTIECR | 0.25 | 0.25 | 0.25 | 0.50 | 1 | 2.25 | SC-CIS / FC-CIS | 0.47 |
| Q9UMF0 | ICAM5 | Intercellular adhesion molecule 5 | TFSLSPDAPR | 1 | 1 | 1 | 0.50 | 1.0 | 4.50 | CTRL / RRMS | 0.12 |
| F5GWP8 | JUP | Junction plakoglobin | LLNDEDPVVVTK | 0.25 | 1 | 1 | 0.50 | 1.0 | 3.75 | CTRL / RRMS | 0.21 |
| P14923 | JUP | Junction plakoglobin | LLNQPNQWPLVK | 0.25 | 1 | 1 | 0.50 | 1.0 | 3.75 | CTRL / RRMS | 0.45 |
| Q9BY79-2 | MFRP | Membrane frizzled-related protein | LELSPEPEGPLLR | 0.25 | 0.25 | 0.25 | 0.50 | 1 | 2.25 | SC-CIS / FC-CIS | 7.07 |
| H3BMA1 | MSLN | Mesothelin | TDAVLPLTVAEVQK | 0 | 1 | 1 | 0.50 | 1.0 | 3.50 | CTRL / RRMS | 2.43 |
| Q5VU43-11 | PDE4DIP | Myomegalin | NWEDVPGDQVK | 0.25 | 1 | 1 | 0.50 | 1.0 | 3.75 | CTRL / RRMS | 0.31 |

| | | | | | | | | | | | |
|---|---|---|---|---|---|---|---|---|---|---|---|
| P34059 | GALNS | N-acetylgalactosamine-6-sulfatase | FPLSFASAEYQEALSR | 0.25 | 0.25 | 0.25 | 0.50 | 1 | 2.25 | SC-CIS / FC-CIS | 0.19 |
| P16519-2 | PCSK2 | Neuroendocrine convertase 2 | EELEEELDEAVER | 0 | 1 | 1 | 0.50 | 1.0 | 3.50 | CTRL / RRMS | 4.45 |
| O15240 | VGF | Neurosecretory protein VGF | QQETAAAETETR | 0.25 | 0.25 | 0.25 | 0.50 | 1 | 2.25 | SC-CIS / FC-CIS | 3.86 |
| O15240 | VGF | Neurosecretory protein VGF | SQTHSLPAPESPEPAAPPR | 0.25 | 0.25 | 0.25 | 0.50 | 1 | 2.25 | SC-CIS / FC-CIS | 2.65 |
| Q14697 | GANAB | Neutral alpha-glucosidase AB | LSFQHDPETSVLVLR | 0.25 | 0.25 | 0.25 | 0.50 | 1 | 2.25 | SC-CIS / FC-CIS | 1.44 |
| Q5VST9 | OBSCN | Obscurin | PAPVTFPTAR | 0.25 | 0.25 | 0.25 | 0.50 | 1 | 2.25 | SC-CIS / FC-CIS | 1.84 |
| Q96BZ4 | PLD4 | Phospholipase D4 | YWPVLDNALR | 0.25 | 0.25 | 0.25 | 0.50 | 1 | 2.25 | SC-CIS / FC-CIS | 4.68 |
| F5H827 | POMGNT1 | Protein O-linked-mannose beta-1,2-N-acetylglucosaminyltransferase 1 | LLSEAEVLDHSK | 0.25 | 0.25 | 0.25 | 0.50 | 1 | 2.25 | SC-CIS / FC-CIS | 1.92 |
| P00734 | F2 | Prothrombin | VTGWGNLK | 0.25 | 0.25 | 0.25 | 0.50 | 1 | 2.25 | SC-CIS / FC-CIS | 3.31 |
| P78509-3 | RELN | Reelin | RQLITSFLDSSQSR | 0.25 | 0.25 | 0.25 | 0.50 | 1 | 2.25 | SC-CIS / FC-CIS | 2.85 |
| P13521 | SCG2 | Secretogranin-2 | VLEYLNQEK | 0.25 | 0.25 | 0.25 | 0.50 | 1 | 2.25 | SC-CIS / FC-CIS | 2.48 |
| Q6ZRP7 | QSOX2 | Sulfhydryl oxidase 2 | NQAVCHDYDIHFYPTFR | 0.25 | 0.25 | 0.25 | 0.50 | 1 | 2.25 | SC-CIS / FC-CIS | 0.27 |
| E9PHH3 | SDC1 | Syndecan-1 | EGEAVVLPEVEPGLTAR | 0.25 | 1 | 1 | 0.50 | 1.0 | 3.75 | CTRL / RRMS | 3.50 |

| | | | | | | | | | | | |
|---|---|---|---|---|---|---|---|---|---|---|---|
| O43493 | TGOLN2 | Trans-Golgi network integral membrane protein 2 | SFLPQVLTEWYIPLEK | 1 | 1 | 1 | 0.50 | 1.0 | 4.50 | CTRL / RRMS | 4.40 |
| P78324 | SIRPA | Tyrosine-protein phosphatase non-receptor type substrate 1 | SGAGTELSVR | 0.25 | 0.25 | 0.25 | 0.50 | 1 | 2.25 | SC-CIS / FC-CIS | 2.26 |
| Q8TAG5 | VSTM2A | V-set and transmembrane domain-containing protein 2A | MQAFEASPMWLQDMK | 0.25 | 0.25 | 0.25 | 0.50 | 1 | 2.25 | SC-CIS / FC-CIS | 2.85 |
| P54289-2 | CACNA2D1 | Voltage-dependent calcium channel subunit alpha-2/delta-1 | PMVLAGDK | 0.25 | 0.25 | 0.25 | 0.50 | 1 | 2.25 | SC-CIS / FC-CIS | 2.10 |
| P54289-2 | CACNA2D1 | Voltage-dependent calcium channel subunit alpha-2/delta-1 | RRPWYIQGAASPK | 0.25 | 0.25 | 0.25 | 0.50 | 1 | 2.25 | SC-CIS / FC-CIS | 2.22 |

The results of peptide quantification using Maxquant software and of statistical analysis based on the five tests used to select significant peptides in the two comparisons (CTRL *vs.* RRMS and FC-CIS *vs.* SC-CIS) are indicated. Scores from individual tests (0, 0.25, 0.5, 1) and global scores (0 to 4.5) were calculated for each peptide as described in the Materials and Methods section. Fold changes between the groups were expressed ratios of peptide relative intensities.

### Supplementary Table 3: Proteins differentially secreted by OPCs upon TNFα or sFasL exposure.

| Protein ID | Gene name | Protein name | Number of Peptides | # significant | Median ratio | SecP NN-score | SignalP | Treatment | Identified in Human CSF |
|---|---|---|---|---|---|---|---|---|---|
| P62425 | Rpl7a | 60S ribosomal protein L7a | 13 | 2 | 0.159 | 0.423 | - | TNFα | |
| Q80ZA3 | Serpinf1 | Alpha-2 antiplasmin | 18 | 2 | -1.143 | 0.803 | + | TNFα | + |
| Q80ZA3 | Serpinf1 | Alpha-2 antiplasmin | 18 | 3 | -1.266 | 0.803 | + | sFasL | + |
| P06238 | A2m | Alpha-2-macroglobulin | 77 | 3 | -1.309 | 0.552 | + | TNFα | |
| P06238 | A2m | Alpha-2-macroglobulin | 77 | 2 | -1.393 | 0.552 | + | sFasL | |
| B1WC49 | Api5 | Api5 protein | 11 | 2 | 1.106 | 0.244 | - | TNFα | |
| B1WC49 | Api5 | Api5 protein | 11 | 3 | 1.248 | 0.244 | - | sFasL | |
| D4A4M3 | Atrip | ATR-interacting protein | 2 | 3 | -5.087 | 0.333 | - | TNFα | |
| Q924T4 | Lfng | Beta-1,3-N-acetylglucosaminyltransferase lunatic fringe | 5 | 2 | -0.883 | 0.814 | + | TNFα | + |
| Q3ZB98 | Bcas1 | Breast carcinoma-amplified sequence 1 homolog (Fragment) | 6 | 2 | -2.435 | 0.140 | - | sFasL | |
| Q63120 | Abcc2 | Canalicular multispecific organic anion transporter 1 | 1 | 2 | 2.223 | 0.593 | - | TNFα | |
| Q9R1T3 | Ctsz | Cathepsin Z | 16 | 2 | -0.730 | 0.798 | + | sFasL | |

| | | | | | | | | | |
|---|---|---|---|---|---|---|---|---|---|
| B0BNA5 | Cotl1 | Coactosin-like protein | 13 | 2 | -0.707 | 0.808 | - | TNFα | + |
| P31720 | C1qa | Complement C1q subcomponent subunit A | 11 | 2 | 0.559 | 0.623 | + | sFasL | |
| P31721 | C1qb | Complement C1q subcomponent subunit B | 8 | 2 | 0.284 | 0.908 | + | TNFα | |
| P31721 | C1qb | Complement C1q subcomponent subunit B | 8 | 2 | 0.711 | 0.908 | + | sFasL | |
| P31722 | C1qc | Complement C1q subcomponent subunit C | 7 | 2 | 1.0646 | 0.737 | + | sFasL | |
| M0RBF1 | C3 | Complement C3 | 104 | 3 | 1.643 | 0.570 | + | TNFα | |
| D3ZD97 | Dhx15 | DEAH (Asp-Glu-Ala-His) box polypeptide 15 (Predicted), isoform CRA_b | 15 | 2 | 1.285 | 0.369 | - | TNFα | |
| P47942 | Dpysl2 | Dihydropyrimidinase-related protein 2 | 41 | 2 | 0.835 | 0.416 | - | TNFα | |
| P80067 | Ctsc | Dipeptidyl peptidase 1 | 8 | 2 | -0.875 | 0.735 | + | sFasL | + |
| Q14TE9 | Top2b | DNA topoisomerase 2 | 4 | 2 | 1.554 | 0.118 | - | TNFα | |
| Q14TE9 | Top2b | DNA topoisomerase 2 | 4 | 2 | 1.783 | 0.118 | - | sFasL | |
| D4A599 | Dnahc17 | Dynein, axonemal, heavy chain 17 | 2 | 2 | -2.167 | ND | | TNFα | |
| P02793 | Ftl1 | Ferritin light chain 1 | 14 | 2 | -0.763 | 0.446 | - | TNFα | + |
| Q08326 | Rabif | Guanine nucleotide exchange factor MSS4 | 1 | 2 | -1.862 | 0.353 | - | TNFα | |

| | | | | | | | | | |
|---|---|---|---|---|---|---|---|---|---|
| P08025 | Igf1 | Insulin-like growth factor I | 2 | 2 | -1.531 | 0.926 | + | sFasL | |
| P12843 | Igfbp2 | Insulin-like growth factor-binding protein 2 | 12 | 3 | -1.402 | 0.788 | + | TNFα | + |
| P15473 | Igfbp3 | Insulin-like growth factor-binding protein 3 | 8 | 2 | -1.907 | 0.793 | + | TNFα | + |
| P15473 | Igfbp3 | Insulin-like growth factor-binding protein 3 | 8 | 2 | -1.385 | 0.793 | + | sFasL | + |
| P24594 | Igfbp5 | Insulin-like growth factor-binding protein 5 | 10 | 4 | -3.460 | 0.926 | + | TNFα | |
| P24594 | Igfbp5 | Insulin-like growth factor-binding protein 5 | 10 | 3 | -4.168 | 0.926 | + | sFasL | |
| Q7TP98 | Ilf2 | Interleukin enhancer-binding factor 2 | 15 | 2 | 0.923 | 0.487 | - | TNFα | |
| Q7TP98 | Ilf2 | Interleukin enhancer-binding factor 2 | 15 | 2 | 0.673 | 0.487 | - | sFasL | |
| G3V7Q7 | Iqgap1 | IQ motif containing GTPase activating protein 1 (Predicted), isoform CRA_b | 13 | 2 | 2.245 | 0.248 | - | sFasL | + |
| Q9JIL3-2 | Ilf3 | Isoform 2 of Interleukin enhancer-binding factor 3 | 17 | 2 | 0.792 | 0.058 | - | sFasL | |
| P27471 | Klrb1a | Killer cell lectin-like receptor subfamily B member 1A | 1 | 2 | -2.063 | 0.894 | - | TNFα | |
| D3ZN61 | Lgi3 | Leucine-rich repeat LGI family, member 3 (Predicted), isoform CRA_b | 22 | 2 | -0.966 | 0.719 | + | sFasL | |
| G3V7A5 | Ldlr | Low density lipoprotein receptor, isoform CRA_a | 20 | 3 | -1.155 | 0.471 | + | TNFα | + |

| | | | | | | | | | |
|---|---|---|---|---|---|---|---|---|---|
| P00697 | Lyz1 | Lysozyme C-1 | 8 | 2 | 1.223 | 0.907 | + | TNFα | + |
| Q63691 | Cd14 | Monocyte differentiation antigen CD14 | 10 | 2 | 0.949 | 0.731 | + | TNFα | + |
| Q6AY12 | Cyb5r2 | NADH-cytochrome b5 reductase 2 | 2 | 2 | -1.888 | 0.612 | - | TNFα | |
| P30152 | Lcn2 | Neutrophil gelatinase-associated lipocalin | 5 | 3 | 1.896 | 0.801 | + | TNFα | + |
| P13084 | Npm1 | Nucleophosmin | 8 | 2 | 0.651 | 0.803 | - | TNFα | |
| D3ZLH5 | Plxnb3 | Plexin-B3 | 19 | 2 | 0.946 | 0.68 | + | TNFα | + |
| F1MA59 | Col4a1 | Protein Col4a1 | 33 | 3 | -1.886 | 0.043 | + | TNFα | + |
| G3V904 | Pld4 | Protein Pld4 | 14 | 2 | -1.153 | 0.83 | - | sFasL | |
| Q91XU1 | Qki | Protein quaking | 9 | 2 | 1.248 | 0.368 | - | TNFα | |
| F1LVV4 | Rcc2 | Protein Rcc2 (Fragment) | 17 | 2 | 0.849 | 0.45 | - | TNFα | |
| D3ZMS1 | Sf3b2 | Protein Sf3b2 | 5 | 2 | 1.381 | 0.108 | - | TNFα | |
| B2RZ74 | Snrnp70 | Protein Snrnp70 | 6 | 2 | 1.391 | 0.467 | - | sFasL | |
| M0R5U3 | Tatdn3 | Protein Tatdn3 | 5 | 2 | -0.281 | 0.276 | - | TNFα | |
| M0RD63 | Ranbp3 | RAN-binding protein 3 | 10 | 2 | 1.055 | 0.213 | - | TNFα | |
| P12346 | Tf | Serotransferrin | 49 | 2 | 0.924 | 0.617 | + | TNFα | |

| | | | | | | | |
|---|---|---|---|---|---|---|---|
| P32821 | Trypsin V-A | 1 | 2 | -2.384 | 0.864 | + | TNFα |

The number of replicates where a significant protein ratio was found and the median ratio measured for each protein in the three biological replicates are indicated. The NN-score corresponds to the score provided by the SecretomeP 2.0 web server (https://services.healthtech.dtu.dk/services/SecretomeP-2.0/). A score higher than 0.5 indicates a high probability of secretion *via* a non-classical secretion mechanism. The presence of signal peptides in the identified protein sequences was predicted by the SignalP algorithm (https://services.healthtech.dtu.dk/service.php?SignalP-5.0).

**Supplementary Table 4. Peptides designed for the qualification step PRM experiments**

| Peptide Name | Peptide Sequence | Origin | Groups compared |
|---|---|---|---|
| AGT_1 | SLDFTELDVAAEK | Litterature | NA |
| AGT_2 | VLSALQAVQGLLVAQGR | Litterature | NA |
| AGT_3 | LQAILGVPWK | Litterature | NA |
| ALCAM_1 | SSNTYTLTDVR | Litterature | NA |
| ALCAM_2 | ALFLETEQLK | Litterature | NA |
| ALCAM_3 | ESLTLIVEGKPQIK | Litterature | NA |
| APOH_1 | ATFGCHDGYSLDGPEEIECTK | Label-free peptide | FC-CIS vs. SC-CIS |
| APOH_2 | FICPLTGLWPINTLK | Label-free peptide | FC-CIS vs. SC-CIS |
| APOH_3 | *WSPELPVCAPIICPPPSIPTFATLR | Label-free peptide | FC-CIS vs. SC-CIS |
| BASP1_1 | ESEPQAAAEPAEAK | Label-free protein | FC-CIS vs. SC-CIS |
| BASP1_2 | APEQEQAAPGPAAGGEAPK | Label-free protein | FC-CIS vs. SC-CIS |
| BASP1_3 | ETPAATEAPSSTPK | Label-free protein | FC-CIS vs. SC-CIS |
| BSG_1 | FFVSSSQGR | Litterature | NA |
| CACNA2D1_1 | VFTFSVGQHNYDR | Label-free peptide | FC-CIS vs. SC-CIS |
| CACNA2D1_2 | YQDLYTVEPNNAR | Label-free peptide | FC-CIS vs. SC-CIS |
| CACNA2D1_3 | TASGVNQLVDIYEK | Label-free peptide | FC-CIS vs. SC-CIS |
| CADM2_1 | SDDGVAVICR | Label-free peptide | FC-CIS vs. SC-CIS |
| CADM2_2 | TFTVSSTLDFR | Label-free peptide | FC-CIS vs. SC-CIS |
| CADM2_3 | GKPLPEPVLWTK | Label-free peptide | FC-CIS vs. SC-CIS |
| CBLN4_1 | VAFSAVR | Label-free peptide | CTRL vs. RRMS; FC-CIS vs. SC-CIS |

| | | | |
|---|---|---|---|
| CBLN4_2 | GSSSSPLGISVR | Label-free peptide | CTRL vs. RRMS; FC-CIS vs. SC-CIS |
| CBLN4_3 | GIYSFSFHVIK | Label-free peptide | CTRL vs. RRMS; FC-CIS vs. SC-CIS |
| CD14_1 | LTVGAAQVPAQLLVGALR | Cultures | TNF$\alpha$ |
| CD14_2 | *FPAIQNLALR | Cultures | TNF$\alpha$ |
| CD14_3 | AFPALTSLDLSDNPGLGER | Cultures | TNF$\alpha$ |
| CD27_1 | *HCNSGLLVR | Label-free peptide | CTRL vs. RRMS |
| CECR1_1 | LLPVYELSGEHHDEEWSVK | Label-free protein/peptide | CTRL vs. RRMS |
| CECR1_2 | *SQVFNILR | Label-free protein/peptide | CTRL vs. RRMS |
| CECR1_3 | LPYFFHAGETDWQGTSIDR | Label-free protein/peptide | CTRL vs. RRMS |
| CFHR1_1 | CLHPCVISR | Label-free peptide | FC-CIS vs. SC-CIS |
| CFHR1_2 | ITCTEEGWSPTPK | Label-free peptide | FC-CIS vs. SC-CIS |
| CFHR1_3 | INHGILYDEEK | Label-free peptide | FC-CIS vs. SC-CIS |
| CHGA_1 | SGELEQEEER | Label-free peptide | FC-CIS vs. SC-CIS |
| CHGA_2 | SEALAVDGAGK | Label-free peptide | FC-CIS vs. SC-CIS |
| CHGA_3 | VAHQLQALR | Label-free peptide | FC-CIS vs. SC-CIS |
| CHI3L1_1 | ILGQQVPYATK | Label-free protein/peptide | CTRL vs. RRMS |
| CHI3L1_2 | *EAGTLAYYEICDFLR | Label-free protein/peptide | CTRL vs. RRMS |
| CHI3L1_3 | LVCYYTSWSQYR | Label-free protein/peptide | CTRL vs. RRMS |

| | | | |
|---|---|---|---|
| CHI3L2_1 | *LLLTAGVSAGR | Label-free protein/peptide | CTRL vs. RRMS |
| CHI3L2_2 | *GPSSYYNVEYAVGYWIHK | Label-free protein/peptide | CTRL vs. RRMS |
| CHI3L2_3 | ENTHFTVLIHELAEAFQK | Label-free protein/peptide | CTRL vs. RRMS |
| CHIT1_1 | VGAPATGSGTPGPFTK | Label-free protein/peptide | CTRL vs. RRMS |
| CHIT1_2 | YPLIQTLR | Label-free protein/peptide | CTRL vs. RRMS |
| CHIT1_3 | *DNQWVGFDDVESFK | Label-free protein/peptide | CTRL vs. RRMS |
| CNTN2_1 | VTVTPDGTLIIR | Label-free peptide | CTRL vs. RRMS |
| CNTN2_2 | IIVQAQPEWLK | Label-free peptide | CTRL vs. RRMS |
| CNTN2_3 | *WGCAAAGK | Label-free peptide | CTRL vs. RRMS |
| CNTN3_1 | VLGSPTPLVLR | Litterature | NA |
| CNTN3_2 | IEVQFPETLPAAK | Litterature | NA |
| CNTN3_3 | ITLHCEAR | Litterature | NA |
| COL4A1_1 | HSQTIDDPQCPSGTK | Cultures | TNF$\alpha$ |
| COL4A1_2 | ILYHGYSLLYVQGNER | Cultures | TNF$\alpha$ |
| COL4A1_3 | AHGQDLGTAGSCLR | Cultures | TNF$\alpha$ |
| COTL1_1 | LFAFVR | Cultures | TNF$\alpha$ |
| CRTAC1_1 | GVALADFNR | Label-free peptide | CTRL vs. RRMS |
| CRTAC1_2 | GNQGFNNNWLR | Label-free peptide | CTRL vs. RRMS |

| | | | |
|---|---|---|---|
| CRTAC1_3 | WEDILSDEVNVAR | Label-free peptide | CTRL vs. RRMS |
| CTSC_1 | ILHLPTSWDWR | Cultures | sFasL |
| CTSC_2 | NVHGINFVSPVR | Cultures | sFasL |
| CTSC_3 | GIYHHTGLR | Cultures | sFasL |
| CTSZ_1 | NVDGVNYASITR | Cultures | sFasL |
| CTSZ_2 | VGDYGSLSGR | Cultures | sFasL |
| CTSZ_3 | NSWGEPWGER | Cultures | sFasL |
| DSC1_1 | *VTIFTVPENCR | Label-free protein/peptide | CTRL vs. RRMS; FC-CIS vs. SC-CIS |
| DSC1_2 | ILQQIPDHPK | Label-free protein/peptide | CTRL vs. RRMS; FC-CIS vs. SC-CIS |
| DSC1_3 | IEDDNDNAPYFEHR | Label-free protein/peptide | CTRL vs. RRMS; FC-CIS vs. SC-CIS |
| DSP_1 | *QLQNIIQATSR | Label-free peptide | CTRL vs. RRMS |
| DSP_2 | LTYEIEDEK | Label-free peptide | CTRL vs. RRMS |
| DSP_3 | *LNDSILQATEQR | Label-free peptide | CTRL vs. RRMS |
| EFNB2_1 | LDIICPK | Litterature | NA |
| EFNB2_2 | ENTPLLNCAKPDQDIK | Litterature | NA |
| EPHA4_1 | VYPANEVTLLDSR | Litterature | NA |
| EPHA4_2 | GLNPLTSYVFHVR | Litterature | NA |
| EPHA4_3 | CPPHSYSVWEGATSCTCDR | Litterature | NA |
| F2_1 | SGIECQLWR | Label-free peptide | FC-CIS vs. SC-CIS |
| F2_2 | ETAASLLQAGYK | Label-free peptide | FC-CIS vs. SC-CIS |

| | | | |
|---|---|---|---|
| F2_3 | YGFYTHVFR | Label-free peptide | FC-CIS vs. SC-CIS |
| F5_1 | AVQPGETYTYK | Label-free protein | FC-CIS vs. SC-CIS |
| F5_2 | EDGILGPIIR | Label-free protein | FC-CIS vs. SC-CIS |
| F5_3 | LAAALGIR | Label-free protein | FC-CIS vs. SC-CIS |
| FCGBP_1 | GNPAVSYVR | Label-free peptide | FC-CIS vs. SC-CIS |
| FCGBP_2 | APGWDPLCWDECR | Label-free peptide | FC-CIS vs. SC-CIS |
| FCGBP_3 | SPANCPLSCPANSR | Label-free peptide | FC-CIS vs. SC-CIS |
| FGA_1 | ESSSHHPGIAEFPSR | Label-free peptide | FC-CIS vs. SC-CIS |
| FGA_2 | *GSESGIFTNTK | Label-free peptide | FC-CIS vs. SC-CIS |
| FGA_3 | QFTSSTSYNR | Label-free peptide | FC-CIS vs. SC-CIS |
| FOLR1_1 | EDCEQWWEDCR | Label-free peptide | FC-CIS vs. SC-CIS |
| FOLR1_2 | *DVSYLYR | Label-free peptide | FC-CIS vs. SC-CIS |
| FTL_1 | LNQALLDLHALGSAR | Cultures | TNF$\alpha$ |
| FTL_2 | TDPHLCDFLETHFLDEEVK | Cultures | TNF$\alpha$ |
| FTL_3 | LGGPEAGLGEYLFER | Cultures | TNF$\alpha$ |
| GALNS_1 | YYEEFPINLK | Label-free protein/peptide | FC-CIS vs. SC-CIS |
| GALNS_2 | LPLIFHLGR | Label-free protein/peptide | FC-CIS vs. SC-CIS |
| GALNS_3 | *FPLSFASAEYQEALSR | Label-free protein/peptide | FC-CIS vs. SC-CIS |
| GANAB_1 | FGAVWTGDNTAEWDHLK | Label-free peptide | FC-CIS vs. SC-CIS |
| GANAB_2 | *LSFQHDPETSVLVLR | Label-free peptide | FC-CIS vs. SC-CIS |

| | | | |
|---|---|---|---|
| GANAB_3 | GLLEFEHQR | Label-free peptide | FC-CIS vs. SC-CIS |
| GC_1 | SLGECCDVEDSTTCFNAK | Litterature | NA |
| GC_2 | VPTADLEDVLPLAEDITNILSK | Litterature | NA |
| GC_3 | HLSLLTTLSNR | Litterature | NA |
| GFRA2_1 | DFTENPCLR | Label-free peptide | FC-CIS vs. SC-CIS |
| GFRA2_2 | LADFHANCR | Label-free peptide | FC-CIS vs. SC-CIS |
| GFRA2_3 | *QTILPSCSYEDK | Label-free peptide | FC-CIS vs. SC-CIS |
| GGT7_1 | EQALHWVAETLK | Label-free protein | CTRL vs. RRMS |
| GGT7_2 | IALALASR | Label-free protein | CTRL vs. RRMS |
| GGT7_3 | VDVLSWVHGSR | Label-free protein | CTRL vs. RRMS |
| GNPTG_1 | RDPSPVSGPVHLFR | Litterature | NA |
| GNPTG_2 | TLFEDAGYLK | Litterature | NA |
| GOLIM4_1 | QAELEEGRPQHQEQLR | Label-free peptide | CTRL vs. RRMS |
| GOLIM4_2 | EAANLLEGHAR | Label-free peptide | CTRL vs. RRMS |
| GOLIM4_3 | *NNDVWQNHEAVPGR | Label-free peptide | CTRL vs. RRMS |
| GRIA4_1 | LQNILEQIVSVGK | Label-free peptide | FC-CIS vs. SC-CIS |
| GRIA4_2 | GVFAIFGLYDK | Label-free peptide | FC-CIS vs. SC-CIS |
| GRIA4_3 | *NTDQEYTAFR | Label-free peptide | FC-CIS vs. SC-CIS |
| HSP90B1_1 | *FQSSHHPTDITSLDQYVER | Label-free peptide | CTRL vs. RRMS |
| HSP90B1_2 | ELISNASDALDK | Label-free peptide | CTRL vs. RRMS |
| HSP90B1_3 | SILFVPTSAPR | Label-free peptide | CTRL vs. RRMS |
| ICAM2_1 | QVILTLQPTLVAVGK | Label-free peptide | FC-CIS vs. SC-CIS |
| ICAM2_2 | *SFTIECR | Label-free peptide | FC-CIS vs. SC-CIS |

| | | | |
|---|---|---|---|
| ICAM2_3 | VPTVEPLDSLTLFLFR | Label-free peptide | FC-CIS vs. SC-CIS |
| ICAM5_1 | *TFSLSPDAPR | Label-free peptide | CTRL vs. RRMS |
| ICAM5_2 | SDGGAVLALGLLGPVTR | Label-free peptide | CTRL vs. RRMS |
| ICAM5_3 | ASLTLTLLR | Label-free peptide | CTRL vs. RRMS |
| IGBRO_1 | EVQLVESGGGLVQPGGSLR | Label-free protein | CTRL vs. RRMS |
| IGBRO_2 | AEDTAVYYCAR | Label-free protein | CTRL vs. RRMS |
| IGFBP2_1 | LAACGPPPVAPPAAVAAVAGGAR | Cultures | TNF$\alpha$ |
| IGFBP2_2 | TPCQQELDQVLER | Cultures | TNF$\alpha$ |
| IGFBP2_3 | LIQGAPTIR | Cultures | TNF$\alpha$ |
| IGFBP3_1 | ALAQCAPPPAVCAELVR | Cultures | sFasL |
| IGFBP3_2 | CQPSPDEARPLQALLDGR | Cultures | sFasL |
| IGFBP3_3 | SAGSVESPSVSSTHR | Cultures | sFasL |
| IGHG1_1 | TPEVTCVVVDVSHEDPEVK | Label-free peptide | CTRL vs. RRMS |
| IGHG1_2 | GPSVFPLAPSSK | Label-free peptide | CTRL vs. RRMS |
| IGHG1_3 | FNWYVDGVEVHNAK | Label-free peptide | CTRL vs. RRMS |
| IGKC_1 | *SGTASVVCLLNNFYPR | Label-free protein/peptide | CTRL vs. RRMS |
| IGKC_2 | VDNALQSGNSQESVTEQDSK | Label-free protein/peptide | CTRL vs. RRMS |
| IGKC_3 | VYACEVTHQGLSSPVTK | Label-free protein/peptide | CTRL vs. RRMS |
| IGLC2_1 | *SYSCQVTHEGSTVEK | Label-free peptide | CTRL vs. RRMS |
| IGLC2_2 | *ATLVCLISDFYPGAVTVAWK | Label-free peptide | CTRL vs. RRMS |

| | | | |
|---|---|---|---|
| IGLC2_3 | *AAPSVTLFPPSSEELQANK | Label-free peptide | CTRL vs. RRMS |
| IGLEN_1 | LLIYWASTR | Label-free protein/peptide | CTRL vs. RRMS |
| IGLL5_1 | ANPTVTLFPPSSEELQANK | Litterature | NA |
| IGWAH_1 | *SGTSASLAISGLR | Label-free protein/peptide | CTRL vs. RRMS |
| IGWAH_2 | *QSVLTQPPSASGTPGQR | Label-free protein/peptide | CTRL vs. RRMS |
| IQGAP1_1 | LTAEEMDER | Cultures | sFasL |
| JUP_1 | LNYGIPAIVK | Label-free peptide | CTRL vs. RRMS |
| JUP_2 | *LLNDEDPVVVTK | Label-free peptide | CTRL vs. RRMS |
| JUP_3 | *VSVELTNSLFK | Label-free peptide | CTRL vs. RRMS |
| L1CAM_1 | LVVFPTDDISLK | Litterature | NA |
| L1CAM_2 | LVLSDLHLLTQSQVR | Litterature | NA |
| L1CAM_3 | FQLQATTK | Litterature | NA |
| LCN2_1 | SYPGLTSYLVR | Cultures | TNF$\alpha$ |
| LDLR_1 | CIPQFWR | Cultures | TNF$\alpha$ |
| LFNG_1 | SIHCHLYPDTPWCPR | Cultures | TNF$\alpha$ |
| LGALS7_1 | AVVGDAQYHHFR | Label-free peptide | CTRL vs. RRMS |
| LGALS7_2 | *SSLPEGIRPGTVLR | Label-free peptide | CTRL vs. RRMS |
| LGALS7_3 | GPGVPFQR | Label-free peptide | CTRL vs. RRMS |
| LYZ_1 | WESGYNTR | Cultures | TNF$\alpha$ |
| LYZ_2 | STDYGIFQINSR | Cultures | TNF$\alpha$ |

| | | | |
|---|---|---|---|
| LYZ_3 | AWVAWR | Cultures | TNFα |
| MFRP_1 | SLTSLPCYQHFR | Label-free protein/peptide | FC-CIS vs. SC-CIS |
| MFRP_2 | *LELSPEPEGPLLR | Label-free protein/peptide | FC-CIS vs. SC-CIS |
| MMRN2_1 | EAEPLVDIR | Litterature | NA |
| MSLN_1 | VNAIPFTYEQLDVLK | Label-free peptide | CTRL vs. RRMS |
| MSLN_2 | *TDAVLPLTVAEVQK | Label-free peptide | CTRL vs. RRMS |
| MSLN_3 | LDELYPQGYPESVIQHLGYLFLK | Label-free peptide | CTRL vs. RRMS |
| MXRA8_1 | AYGPLFLR | Litterature | NA |
| MXRA8_2 | GAPALLTCVNR | Litterature | NA |
| MXRA8_3 | LLDLYASGER | Litterature | NA |
| NAGLU_1 | LPRPLPAVPGELTEATPNR | Label-free protein | FC-CIS vs. SC-CIS |
| NAGLU_2 | YDLLDLTR | Label-free protein | FC-CIS vs. SC-CIS |
| NAGLU_3 | LLGPGPAADFSVSVER | Label-free protein | FC-CIS vs. SC-CIS |
| NCAM2_1 | LTIYNANIEDAGIYR | Litterature | NA |
| NCAM2_2 | IEIFQTLPVR | Litterature | NA |
| NCAM2_3 | QGEDAEVVCR | Litterature | NA |
| OBSCN_1 | ANCFTEELTNLQVEEK | Label-free peptide | FC-CIS vs. SC-CIS |
| OBSCN_2 | VSFHLHITEPK | Label-free peptide | FC-CIS vs. SC-CIS |
| PCDH17_1 | DDHGLFGLDVK | Label-free protein | CTRL vs. RRMS |
| PCDH17_2 | LEENYDNFYTVVTDRPLDR | Label-free protein | CTRL vs. RRMS |
| PCDH17_3 | NAGLGYLVSTVR | Label-free protein | CTRL vs. RRMS |

| | | | |
|---|---|---|---|
| PCSK2_1 | *EELEEELDEAVER | Label-free peptide | CTRL vs. RRMS |
| PCSK2_2 | YTDDWFNSHGTR | Label-free peptide | CTRL vs. RRMS |
| PCSK2_3 | FHCVGGSVQDPEK | Label-free peptide | CTRL vs. RRMS |
| PDE4DIP_1 | IYFLEER | Label-free peptide | CTRL vs. RRMS |
| PLD4_1 | TSTDLQVLAAR | Label-free peptide | FC-CIS vs. SC-CIS |
| PLD4_2 | *YWPVLDNALR | Label-free peptide | FC-CIS vs. SC-CIS |
| PLXNB3_1 | AQELVACGQVR | Cultures | TNF$\alpha$ |
| PLXNB3_2 | ELPVPIYVTQGEAQR | Cultures | TNF$\alpha$ |
| POMGNT1_1 | SLGSQAGPALGWR | Label-free peptide | FC-CIS vs. SC-CIS |
| POMGNT1_2 | EAYEVEVHR | Label-free peptide | FC-CIS vs. SC-CIS |
| POMGNT1_3 | *LLSEAEVLDHSK | Label-free peptide | FC-CIS vs. SC-CIS |
| QSOX2_1 | *NQAVCHDYDIHFYPTFR | Label-free peptide | FC-CIS vs. SC-CIS |
| RELN_1 | ITIPLPNAALTR | Label-free peptide | FC-CIS vs. SC-CIS |
| RELN_2 | VIVLLPQK | Label-free peptide | FC-CIS vs. SC-CIS |
| RELN_3 | VPSLVSVVINPELQTPATK | Label-free peptide | FC-CIS vs. SC-CIS |
| SCG2_1 | VPGQGSSEDDLQEEEQIEQAIK | Label-free peptide | FC-CIS vs. SC-CIS |
| SCG2_2 | ALEYIENLR | Label-free peptide | FC-CIS vs. SC-CIS |
| SCG2_3 | IILEALR | Label-free peptide | FC-CIS vs. SC-CIS |
| SDC1_1 | *EGEAVVLPEVEPGLTAR | Label-free peptide | CTRL vs. RRMS |
| SERPINA3_1 | NLAVSQVVHK | Label-free protein/peptide | CTRL vs. RRMS |
| SERPINA3_2 | EQLSLLDR | Label-free protein/peptide | CTRL vs. RRMS |

| | | | |
|---|---|---|---|
| SERPINA3_3 | *LINDYVK | Label-free protein/peptide | CTRL vs. RRMS |
| SERPINF1_1 | LAAAVSNFGYDLYR | Cultures | sFasL; TNFα |
| SERPINF1_2 | TSLEDFYLDEER | Cultures | sFasL; TNFα |
| SERPINF1_3 | DTDTGALLFIGK | Cultures | sFasL; TNFα |
| SIRPA_1 | ATPQHTVSFTCESHGFSPR | Label-free peptide | FC-CIS vs. SC-CIS |
| SIRPA_2 | TETASTVTENK | Label-free peptide | FC-CIS vs. SC-CIS |
| SIRPA_3 | *SGAGTELSVR | Label-free peptide | FC-CIS vs. SC-CIS |
| SLC5A5_1 | EEVAILDDNLVK | Label-free protein | FC-CIS vs. SC-CIS |
| SLC5A5_2 | KPPGFLPTNEDR | Label-free protein | FC-CIS vs. SC-CIS |
| SLC5A5_3 | LFFLGQK | Label-free protein | FC-CIS vs. SC-CIS |
| SPP1_1 | GDSVVYGLR | Litterature | NA |
| SPP1_2 | YPDAVATWLNPDPSQK | Litterature | NA |
| SPP1_3 | QETLPSK | Litterature | NA |
| TGOLN2_1 | EAEDDDTGPEEGSPPK | Label-free peptide | CTRL vs. RRMS |
| TGOLN2_2 | SGAEAQTPEDSPNR | Label-free peptide | CTRL vs. RRMS |
| TGOLN2_3 | SSAEAQTPEDTPNK | Label-free peptide | CTRL vs. RRMS |
| TNXB_1 | FDSFTVQYK | Litterature | NA |
| TNXB_2 | ILISGLEPSTPYR | Litterature | NA |
| TNXB_3 | DAQGQPQAVPVSGDLR | Litterature | NA |
| VGF_1 | EPVAGDAVPGPK | Label-free peptide | FC-CIS vs. SC-CIS |
| VGF_2 | ALAAVLLQALDR | Label-free peptide | FC-CIS vs. SC-CIS |
| VGF_3 | *QQETAAAETETR | Label-free peptide | FC-CIS vs. SC-CIS |

| | | | |
|---|---|---|---|
| VSTM2A_1 | VTDANYGELQEHK | Label-free peptide | FC-CIS vs. SC-CIS |
| VSTM2A_2 | FTEFPR | Label-free peptide | FC-CIS vs. SC-CIS |
| VSTM2A_3 | DEGLYECR | Label-free peptide | FC-CIS vs. SC-CIS |

* indicates peptides significantly up or down regulated in the discovery step. CTRL: symptomatic controls; CIS: clinically isolated syndrome; FC-CIS: fast conversion to MS (< 1year) after a CIS; SC-CIS: slow conversion to MS (>2 years) after CIS; RRMS: relapsing remitting multiple sclerosis patients; sFasL: soluble Fas ligand; TNFα: tumor necrosis factor-alpha.

**Supplementary Table 5: Fold changes of peptides selected for the verification step.**

| Protein Name | Peptide | Peptide Sequence | RRMS *vs.* CTRL | FC-CIS *vs.* SC-CIS | RRMS *vs.* PPMS | RRMS *vs.* INDC |
|---|---|---|---|---|---|---|
| Adenosine deaminase CECR1 | CECR1_1 | LLPVYELSGEHHDEEWSV(K) | 3.59 ** | 0.98 | 2.17 * | 1.69 |
| Adenosine deaminase CECR1 | CECR1_2 | SQVFNIL(R) | 1.81 | 1.11 | 1.27 | 1.35 |
| CD27 antigen | CD27_1 | HCNSGLLV(R) | 22.47 *** | 0.67 | 2.70 * | 6.15 *** |
| Chitinase-3-like protein 1 | CHI3L1_1 | ILGQQVPYAT(K) | 1.88 * | 1.10 | 0.98 | 1.19 |
| Chitinase-3-like protein 1 | CHI3L1_3 | LVCYYTSWSQY(R) | 1.96 * | 0.80 | 0.95 | 1.20 |
| Chitinase-3-like protein 2 | CH3L2_1 | LLLTAGVSAG(R) | 2.37 * | 1.01 | 1.11 | 1.51 |
| Chitinase-3-like protein 2 | CH3L2_2 | GPSSYYNVEYAVGYWIH(K) | 2.97 * | 1.15 | 1.16 | 1.53 |
| Chitotriosidase-1 | CHIT1_1 | VGAPATGSGTPGPFT(K) | 4.54 * | 0.90 | 2.38 | 1.64 |
| Chitotriosidase-1 | CHIT1_3 | DNQWVGFDDVESF(K) | 6.41 ** | 0.92 | 2.44 | 1.86 |
| Complement factor H-related protein 1 | FHR1_3 | INHGILYDEE(K) | 0.56 * | 0.89 | 0.65 | 0.63 |
| Ig kappa chain C region | IGKC_1 | SGTASVVCLLNNFYP(R) | 3.56 * | 0.95 | 2.13 | 3.60 * |
| Ig kappa chain C region | IGKC_2 | VDNALQSGNSQESVTEQDS(K) | 3.81 ** | 1.16 | 2.33 | 3.84 ** |
| Lysozyme C | LYZ_1 | WESGYNT(R) | 1.92 * | 0.95 | 1.28 | 0.93 |
| Neutrophil gelatinase-associated lipocalin | NGAL_1 | SYPGLTSYLV(R) | 0.77 | 0.88 | 0.74 | 0.67 * |
| Reelin | RELN_2 | VIVLLPQ(K) | 0.51 * | 1.28 | 0.86 | 0.69 |
| Syndecan-1 | SDC1_1 | EGEAVVLPEVEPGLTA(R) | 2.59 ** | 1.46 | 1.49 | 2.16 * |

Fold changes of 16 peptides selected for the verification step among the different groups compared. Fold change significance (t-test, p-values) were measured using Msstat in Skyline. * : p-value < 0.05 ; ** : p-value < 0.01 ; *** : p-value < 0.001.

**Supplementary Table 6: Sensitivity and specificity of CSF biomarkers discriminating the different groups.**

| Groups | RRMS, PPMS, SC-CIS, FC-CIS, INDC, NINDC, PINDC | CTRL, ION | Univariate analysis | | Multivariate analysis | |
|---|---|---|---|---|---|---|
| Peptides | n=113 | n=45 | p-value | AUC IC[95%] | p-value | AUC IC[95%] |
| **CHI3L1_y9** | 8592962 [5030770 ; 13774302] | 4191578 [3127934 ; 5512445] | <.0001 | 0.78 [0.7113;0.8552] | **0.0004** | |
| **CHIT1_y12** | 112198 [26058 ; 317992] | 11420 [3458 ; 35183] | 0.0002 | 0.78 [0.7056;0.8501] | **0.0017** | **0.88 [0.8395;0.9403]** |
| **SDC1_y11** | 303486 [196101 ; 578638] | 191394 [129668 ; 276810] | 0.0001 | 0.72 [0.639;0.7974] | **0.0015** | |
| CHI3L2_y8 | 2234839 [1384462 ; 3328136] | 1031757 [789128 ; 1550651] | <.0001 | 0.79 [0.719;0.87] | 0.8434 | |
| CECR1_y15 | 379874 [246855 ; 500311] | 184716 [98283 ; 250500] | <.0001 | 0.81 [0.7425;0.8841] | 0.6776 | |
| IGKC_y8 | 12371120 [6555876 ; 22910822] | 5923137 [4148153 ; 8443602] | 0.0001 | 0.76 [0.6811;0.8292] | 0.9883 | |
| CD27_y7 | 284420 [78242 ; 627524] | 42392 [26661 ; 73064] | 0.0001 | 0.81 [0.747;0.8774] | 0.0685 | |
| NGAL_y8 | 385289 [277589 ; 458851] | 324574 [257587 ; 414018] | 0.0838 | 0.59 [0.4963;0.6888] | 0.8721 | |
| **Groups** | **RRMS, PPMS, SC-CIS, FC-CIS, INDC** | **NINDC, PINDC** | **Univariate analysis** | | **Multivariate analysis** | |

| Peptides | n=87 | n=26 | p-value | AUC IC[95%] | p-value | AUC IC[95%] |
|---|---|---|---|---|---|---|
| **CD27_y7** | 407173 [175665 ; 816668] | 71182.5 [33473 ; 88932] | 0.0003 | 0.87 [0.8078;0.9367] | **0.0017** | **0.91 [0.8564;0.9624]** |
| **SDC1_y11** | 394259 [243206 ; 628153] | 184977.5 [107682 ; 270734] | 0.0002 | 0.82 [0.7393;0.8964] | **0.0297** | |
| CECR1_y15 | 412888 [287770 ; 540565] | 254205 [181573 ; 383306] | 0.0111 | 0.69 [0.575;0.7999] | 0.1854 | |
| CHI3L2_y8 | 2306165 [1588109 ; 3701562] | 1585401 [1134351 ; 2599180] | 0.0438 | 0.65 [0.5359;0.7682] | 0.1075 | |
| CHIT1_y12 | 169795 [17759 ; 402678] | 57467 [30476 ; 155220] | 0.0605 | 0.61 [0.4978;0.7197] | 0.8436 | |
| IGKC_y8 | 14297280 [8135029 ; 27946616] | 5590009 [3572260 ; 11047180] | 0.0012 | 0.81 [0.7163;0.8947] | 0.9064 | |
| CHI3L1_y9 | 8592962 [5030770 ; 13774302] | 8541062 [4555839 ; 14396944] | 0.7702 | 0.51 [0.3814;0.6372] | | |
| NGAL_y8 | 381783 [277589 ; 457131] | 411157 [274912 ; 547633] | 0.1683 | 0.57 [0.4333;0.6985] | | |

| **Groups** | **RRMS, PPMS, SC-CIS, FC-CIS** | **INDC** | **Univariate analysis** | | **Multivariate analysis** | |
|---|---|---|---|---|---|---|
| Peptides | n=74 | n=13 | p-value | AUC IC[95%] | p-value | AUC IC[95%] |
| **SDC1_y11** | 177872 [114147 ; 259174] | 468071 [258136 ; 704891] | **0.0019** | **0.85 [0.7521;0.9547]** | **0.0019** | **0.85 [0.7521;0.9547]** |
| IGKC_y8 | 7774542 [5382441 ; 12654720] | 16595285 [9802823 ; 34762040] | **0.0356** | 0.75 [0.6134;0.8918] | 0.6343 | |
| CD27_y7 | 141177 [103892 ; 318380] | 469457 [260035 ; 946283] | **0.0167** | 0.76 [0.6301;0.8835] | 0.2373 | |

| | | | | | |
|---|---|---|---|---|---|
| NGAL_y8 | 430335 [356360 ; 748360] | 371878 [269925 ; 434625] | **0.0065** | 0.69 [0.4976;0.8787] | 0.2657 |
| CECR1_y15 | 408222 [360991 ; 490899] | 418865.5 [279058 ; 540565] | 0.7936 | 0.52 [0.3624;0.6875] | |
| CHI3L1_y9 | 11937785 [7231543 ; 12922912] | 7976151.5 [4905509 ; 13774302] | 0.2094 | 0.63 [0.4847;0.7814] | |
| CHIT1_y12 | 74945 [5264 ; 131091] | 200208.5 [49496 ; 436242] | 0.1414 | 0.65 [0.492;0.8105] | |
| CHI3L2_y8 | 2229420 [1693552 ; 3607957] | 2332643 [1588109 ; 3701562] | 0.2813 | 0.53 [0.3459;0.7082] | |

| **Groups** | **RRMS** | **PPMS** | **Univariate analysis** | | **Multivariate analysis** | |
|---|---|---|---|---|---|---|
| Peptides | n=30 | n=15 | p-value | AUC IC[95%] | p-value | AUC IC[95%] |
| **CECR1_y15** | 511124.5 [368906 ; 762444] | 253170 [159004 ; 379874] | **0.0042** | **0.85 [0.7364;0.9658]** | **0.0042** | **0.85 [0.7364;0.9658]** |
| NGAL_y8 | 413026 [269792 ; 543017] | 327002 [189399 ; 398146] | **0.048** | 0.68 [0.5299;0.839] | 0.7136 | |
| IGKC_y8 | 23017700 [13986140 ; 50622792] | 12035410 [8135029 ; 31107030] | **0.0956** | 0.70 [0.5289;0.8622] | 0.3944 | |
| CD27_y7 | 692242.5 [380399 ; 1200790] | 265994 [27676 ; 772926] | **0.094** | 0.71 [0.5289;0.8933] | 0.2487 | |
| CHI3L1_y9 | 11198438.5 [6301214 ; 17718706] | 9411357 [4192522 ; 13697746] | 0.196 | 0.61 [0.4305;0.7873] | | |
| CHI3L2_y8 | 3398215 [2231594 ; 4934197] | 1996009 [1163236 ; 3092515] | 0.1181 | 0.69 [0.5295;0.8528] | | |
| CHIT1_y12 | 273658.5 [112198 ; 637876] | 77586 [10180 ; 230341] | 0.5351 | 0.67 [0.4848;0.8486] | | |
| SDC1_y11 | 604636 [250153 ; 814719] | 416163 [302801 ; 673547] | 0.1654 | 0.60 [0.4375;0.7714] | | |

AUCs provided by ROC curves analysis comparing CTRL/ION with MS/INDC/PINDC/NINDC patients; MS/INDC with PINDC/NINDC; MS with INDC; and RRMS with PPMS. Peptides from the univariate analysis with a p-value < 0.1 were used to build a multivariate model. AUCs > 0.8 are in bold. In the multivariate model, significant peptides are in bold and the corresponding AUCs are indicated in grey squares.
