## Supplementary Figures for "Syndecan-1 as specific cerebrospinal fluid biomarker of multiple sclerosis"

| Groups | Diagnosis | Schematic representation of multiple sclerosis related symptoms | MRI | OCB |
|---|---|---|---|---|
| Control, isolated optic neuritis (ION) or other neurological disease patients | CTRL<br>ION<br>INDC<br>PINDC<br>NINDC | 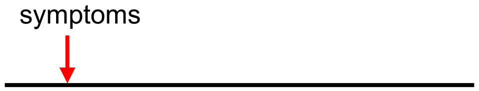 | Normal ? | - / + |
| CIS patients with slow conversion to RRMS (>2 years) | SC-CIS | 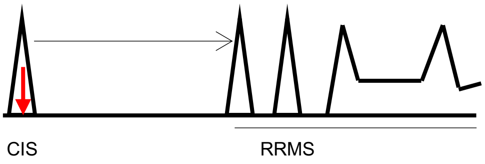 | Swanton | + |
| CIS patients with rapid conversion to RRMS (<1 year) | FC-CIS | 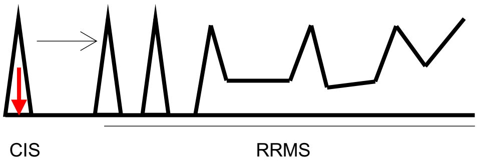 | Swanton | + |
| Relapsing-remitting multiple sclerosis | RRMS | 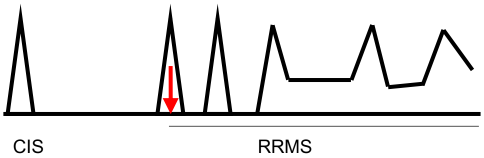 | Swanton | + |
| Primary progressive multiple sclerosis | PPMS | 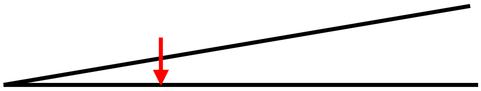 | Swanton | + |

**Supplementary Figure 1**

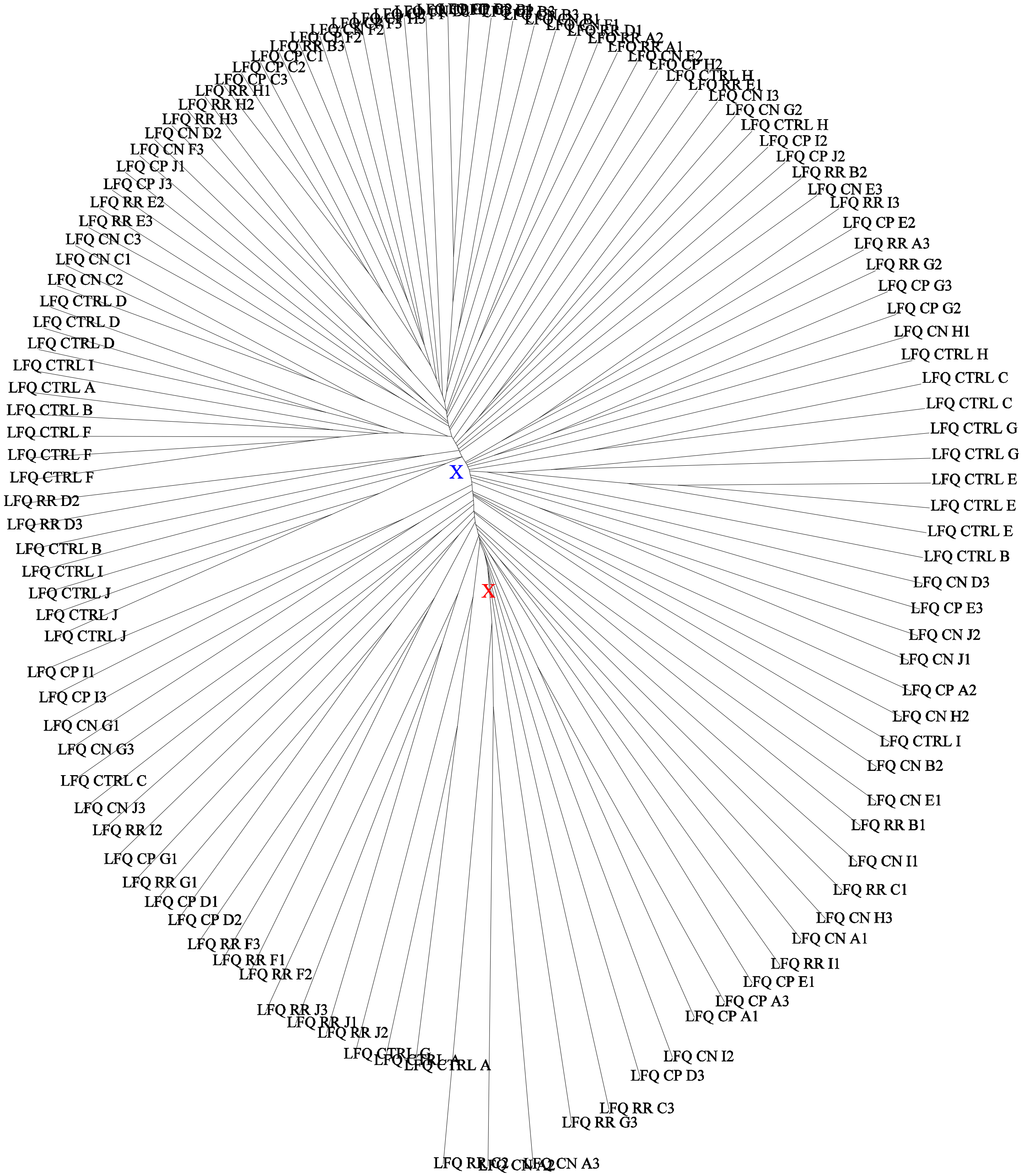


**Supplementary Figure 2**

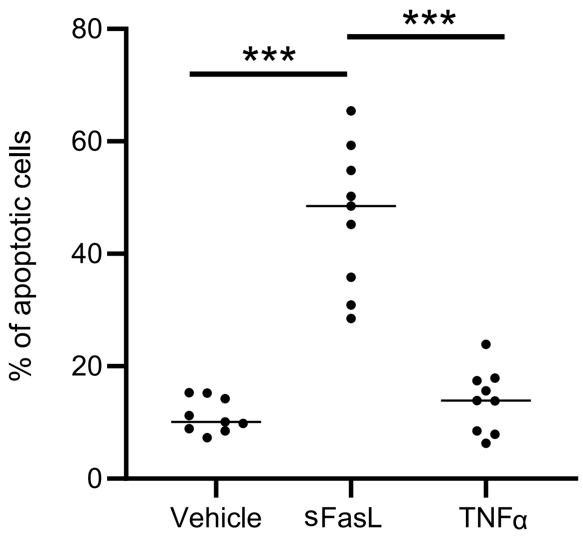


**Supplementary Figure 3**

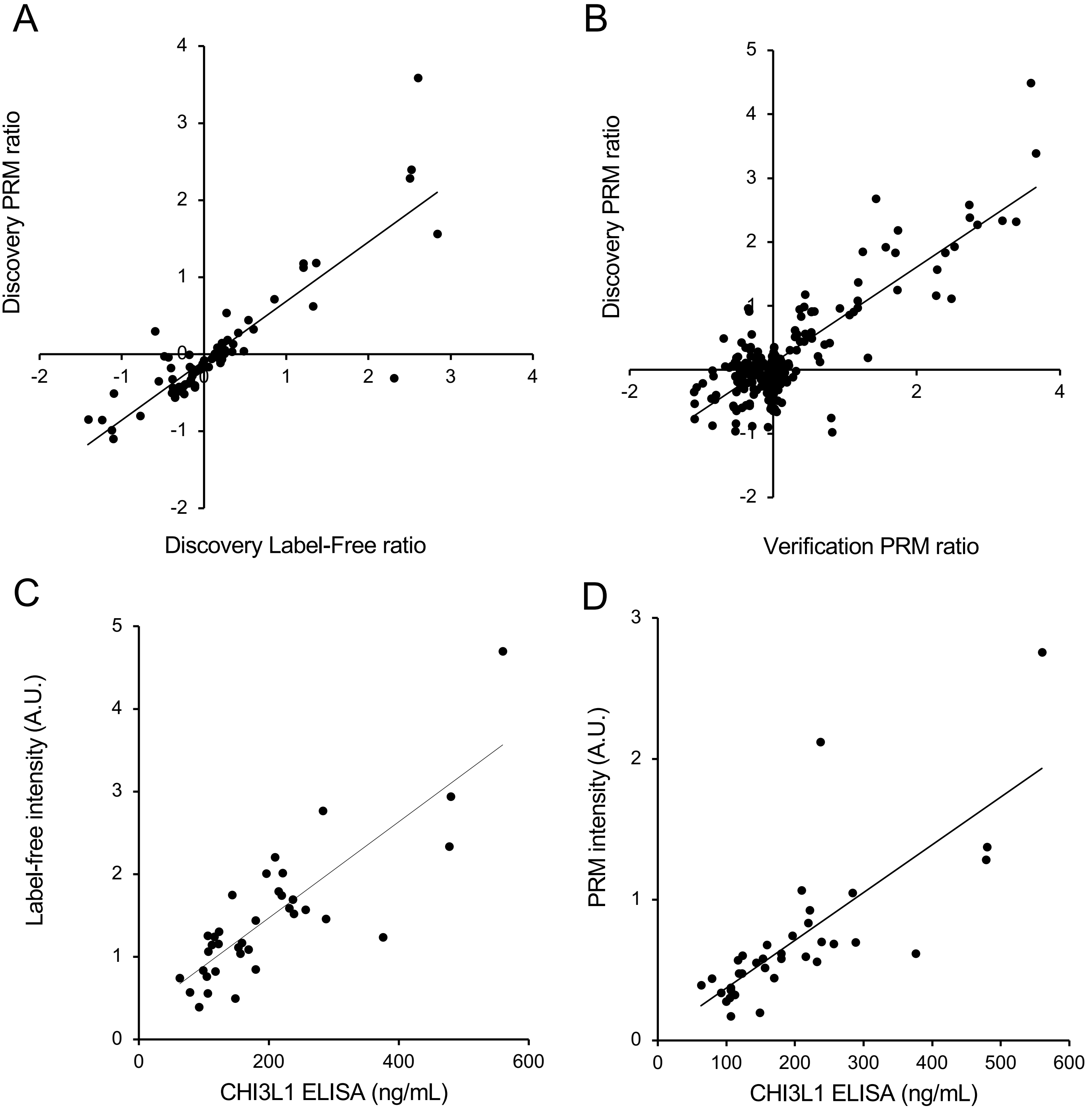


**Supplementary Figure 4**
